# Worldwide phylogeography and history of wheat genetic diversity

**DOI:** 10.1101/477455

**Authors:** François Balfourier, Sophie Bouchet, Sandra Robert, Romain De Oliveira, Hélène Rimbert, Jonathan Kitt, Frédéric Choulet, International Wheat Genome Sequencing Consortium, BreedWheat Consortium, Etienne Paux

## Abstract

Since its domestication in the Fertile Crescent ~8,000 to 10,000 years ago, wheat has undergone a complex history of spread, adaptation and selection. To get better insights into the wheat phylogeography and genetic diversity, we describe allele distribution through time using a set of 4,506 landraces and cultivars originating from 105 different countries genotyped with a high-density SNP array. Although the genetic structure of landraces is collinear to ancient human migration roads, we observe a reshuffling through time, related to breeding programs, with the apparition of new alleles enriched with structural variations that may be the signature of introgressions from wild relatives after 1960.

**One Sentence Summary:** A phylogeographical study reveals the complex history of wheat genetic diversity through time and space.

---

Bread wheat (*Triticum aestivum* L.) is an allohexaploid species originating from two successive rounds of hybridization. The second hybridization event is thought to have occurred in the Fertile Crescent during the Neolithic, ~8,000 to 10,000 years ago (1-4). Wheat was firstly domesticated in a part of the Fertile Crescent area, which is today located in southeastern Turkey. Then bread wheat germplasm has evolved along ancient human migration roads. It has been spread by the first farmers from this area both westwards, to Europe, and eastwards, to Asia, from 8,500 to 2,300 BP (*5*). After their dissemination in Europe and Asia, domesticated wheat populations have slowly adapted to local environments, becoming so-called landraces. From the 16^th^ century, bread wheat was introduced in the New World, first in Latin America then in Australia (5). During the last two centuries, breeding programs were organized to improve these landraces, first in Europe, then in Asia. Finally, after the Second World War, the introduction of dwarf genes in crops during the Green Revolution, in wheat in particular, contributed to dramatic modifications in the genepool over the world (6, 7). Today, with more than 220 million hectares and almost 750 Mt produced every year, wheat is one of the most cultivated and consumed crops worldwide, providing 15% of calories consumed every day (8). Since the transition from hunting-gathering to agriculture, bread wheat has been essential to the rise of civilizations. It has repeatedly been shaped by selection to meet human needs and adaptation to different environments. Here we report on a worldwide phylogeographical study aiming at understanding this complex history of wheat dissemination and differentiation.

Since previous studies highlighted the importance of both geographical and temporal effects in structuring the wheat diversity (9, 10), a set of 4,506 wheat accessions was sampled that represented the worldwide diversity (Data S1) and genotyped using a high-density SNP array containing 280,226 SNPs (11). A set of 113,457 high quality SNPs showing less than 2% missing data was selected (Data S2). The genomic position of these SNPs was determined using the IWGSC RefSeq v1.0 (12). As chip-designed markers can lead to ascertainment bias due to the marker type and selection (13, 14), we inferred haplotype blocks and corresponding alleles along the genome (Figure S1, Data S3). In total, 8,741 haplotypic blocks were identified that presented an allelic frequency distribution close to expectation under Hardy-Weinberg hypothesis compared to bi-allelic markers (Figures S2-S3; Data S3).

In order to have a better understanding of the evolution of allelic diversity through time, we separated accessions in sets that were relevant in terms of agricultural practices: landraces corresponding to the original pool of worldwide diversity, traditional cultivars registered before the Green Revolution and the global introduction of dwarf genes (1960), and modern varieties registered after 1960. When assigning landraces to different genetic groups, we found that eight groups was a good compromise to describe wheat diversity in terms of assignment stability between different runs, geography and migration roads (Figures S4-S7): South Eastern Asia (SEA), Indian Peninsula (INP), Central Asia – Africa (CAA), Caucasus (CAU), South Eastern Europe (SEE), Mediterranean Area (MED), Iberian Peninsula (IBP) and North Western Europe (NWE) (Table S1).

Admixture levels represented by a gradient of colors (Figure 1) reveal the continuous natural selection and differentiation in new environments that occurred during the initial spread of bread wheat from the Fertile Crescent toward Europe and Asia during the Neolithic, along historical roads of migration. These roads include the Danubian road northward through the Balkans, and the Mediterranean westward road across Italy, France, Spain that finally reached UK and Scandinavia, the North African road from Middle East to Africa through Israel and Egypt, and the Silk road from Iran to central Asia, Indian Peninsula, then to China (5). Note that the MED group that is at a crossroad position in the Mediterranean Area is particularly admixed.

**Fig. 1:**
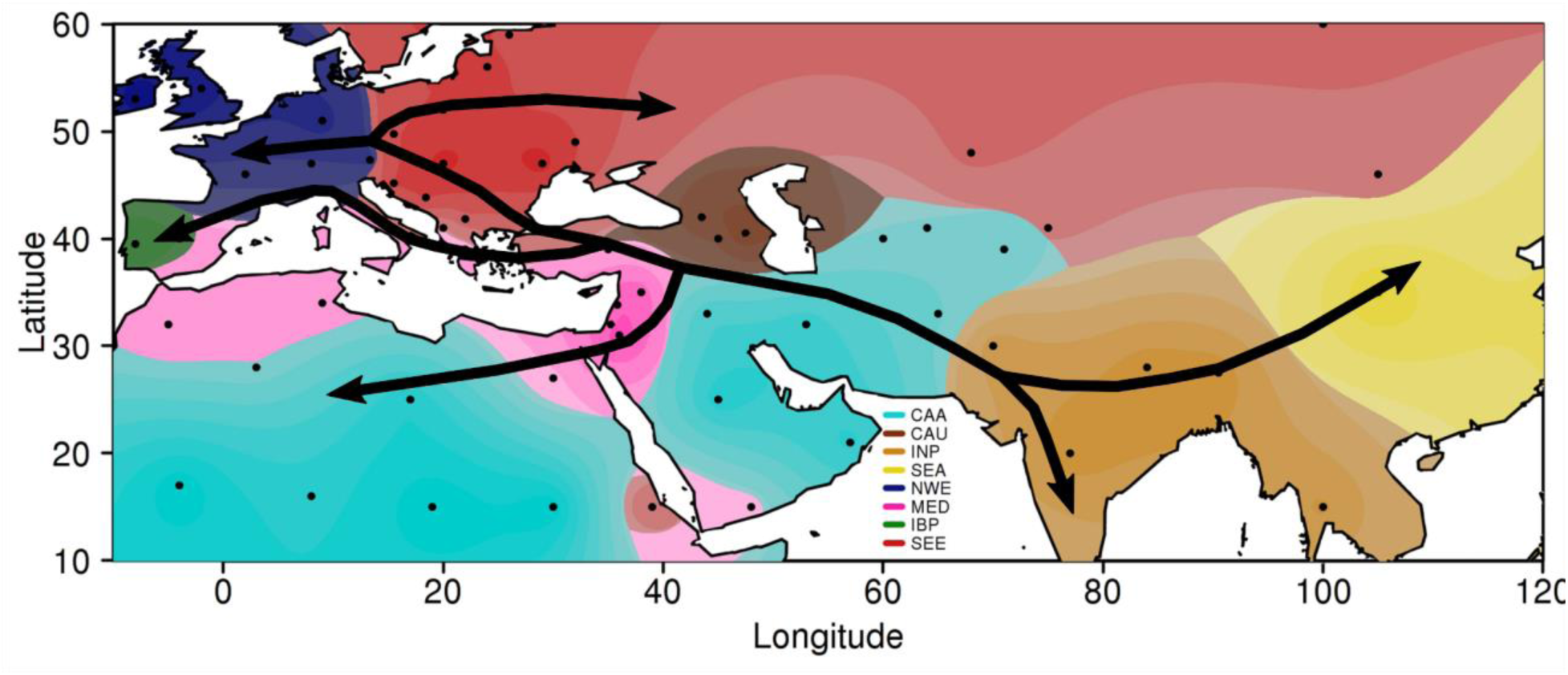
Interpolated map representing the structure of wheat landraces (K=8) and the main human migration roads from the Fertile Crescent. Cyan: CAA, Brown: CAU, Orange: INP, Yellow: SEA, Blue: NEW, Pink: MED, Green: IBP, Red: SEE.

The phylogenetic tree built with 4,403 wheat accessions and the same set of haplotypes revealed three large pools (Figure 2A). The Eastern European & Mediterranean genepool gathered 94% of the SEE, MED and IBP landraces, together with 83% of all traditional and modern varieties not originating from Northern and Western Europe. This is consistent with the successive introductions, first by the Spanish and Portuguese in South America, then by European in the USA, Canada and Australia, and by CIMMYT and ICARDA breeding programs worldwide. Although 94% of Asian landraces (SEA, INP, CAA and CAU) were grouped in the Asian pool, only 7% of the traditional and modern varieties were, while 71% of Asian modern varieties were found within the Eastern European & Mediterranean pool. This shows that Asian breeding programs have mainly relied on European germplasm, and more particularly on Italian one (15). The high differentiation index between modern lines and Asian landraces (SEA, and INP) (Table S5) supports the fact that the original Asian diversity pool did not contribute much to modern varieties. Finally, the Western European genepool contains most of NWE landraces (62 out of 63), as well as 76% of the Northern and Western Europe traditional and modern varieties. This demonstrates that Northern and Western Europe breeding programs have mainly used local landraces. At a finer scale, we defined eleven groups (Figure 2A; Table S3). The distribution of landraces, traditional and modern lines among those groups not only reflects geographical origins but also more recent migrations and variety releases by influent breeding programs (Figures 2B; Supplementary Text; Figures S8-S17).

**Fig. 2:**
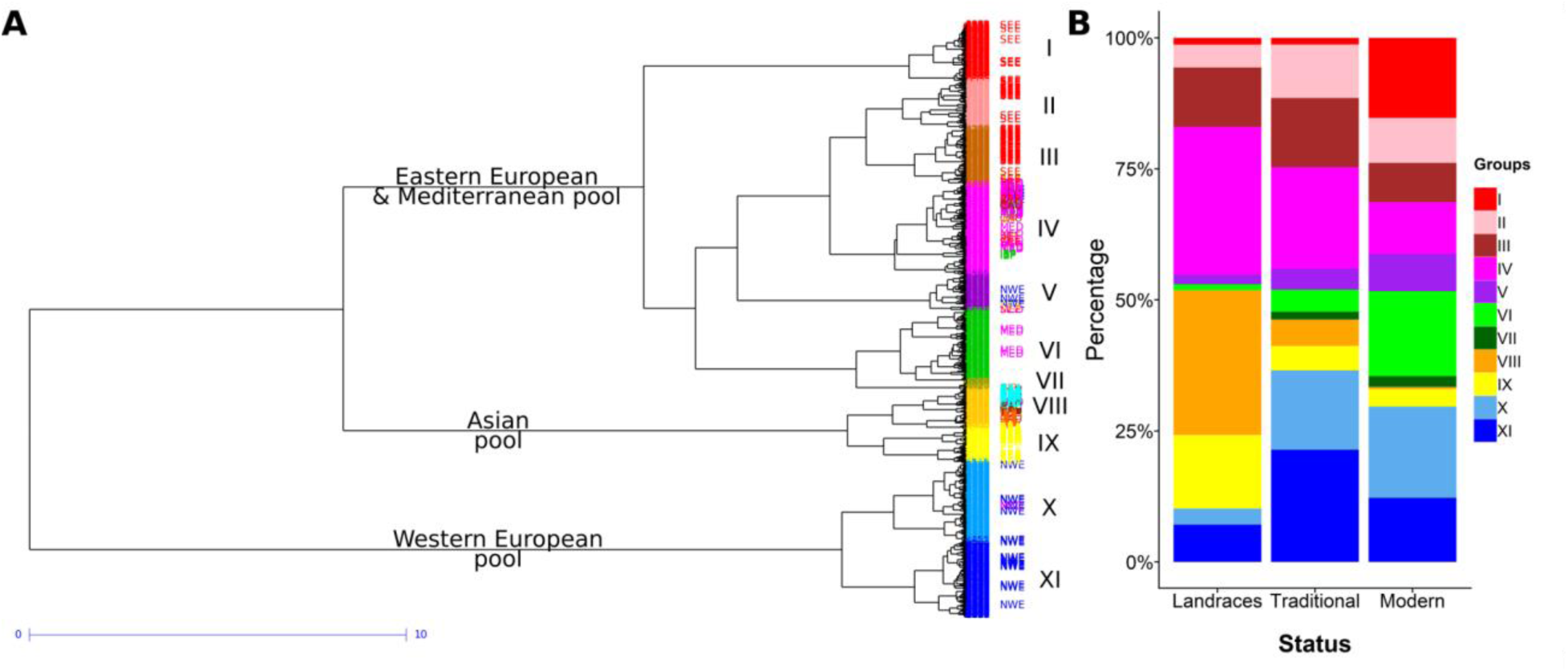
Spatio-temporal structuration of the worldwide diversity. (A) Ward dendrogram showing phylogenetic relationships between 4,403 wheat accessions revealed by 8,741 haplotypes. The different colors correspond to the eleven groups. Landraces are indicated on the right with a color code corresponding to the eight sub-groups. (B) Proportions of the eleven groups in landraces, traditional cultivars and modern varieties.

We observed that the composition in structural variations (SVs) evolved through time. When considering the 14,124 Off-Target Variant (OTV) markers, *i.e.* markers that reveal null alleles and can therefore be used to detect presence / absence variations (16), the number of null alleles appeared to be quite similar in landraces and traditional cultivars (~1,000 OTVs per line). By contrast, we observed a second group of lines with more than 1,700 OTVs, especially in modern varieties registered after 1970 (Figure 3). The structural variations may be due to the use of wild relatives in crop improvement that traces back to the early 1940s and gained prominence during the 1970s and 1980s (17). A detailed analysis of the OTV distribution revealed more than 80 SV blocks greater than 5 Mb with a cumulative length greater than 4,600 Mb, *i.e.* almost one third of the entire bread wheat genome (Table S6). These large blocks likely correspond to alien introgressions, like for example the rye (*Secale cereale*) 1RS on chromosome 1B or the *Aegilops ventricosa* 2NS translocations on chromosome 2A (18, 19). However, while such large blocks can be easily detected, smaller alien segments, referred to as ‘cryptic introgressions’, are more difficult to identify (20). Therefore, it is likely that the total amount of alien DNA present in the current wheat germplasm is by far greater than that. To date, novel alleles have been introgressed from more than 50 species from 13 genera (21), highlighting the importance of these so-called alien introgressions for wheat breeding but also the complex genomic patterns that might have arisen. Although it has been shown that SNP natural genetic diversity in the improved material is significantly lower than the one observed in landraces as a result of domestication and selection (22), breeding programs have brought some new alleles enriched in SVs related to alien introgressions, leading to an overall genetic diversity that is similar between landraces and modern varieties (Figure S18).

**Fig. 3:**
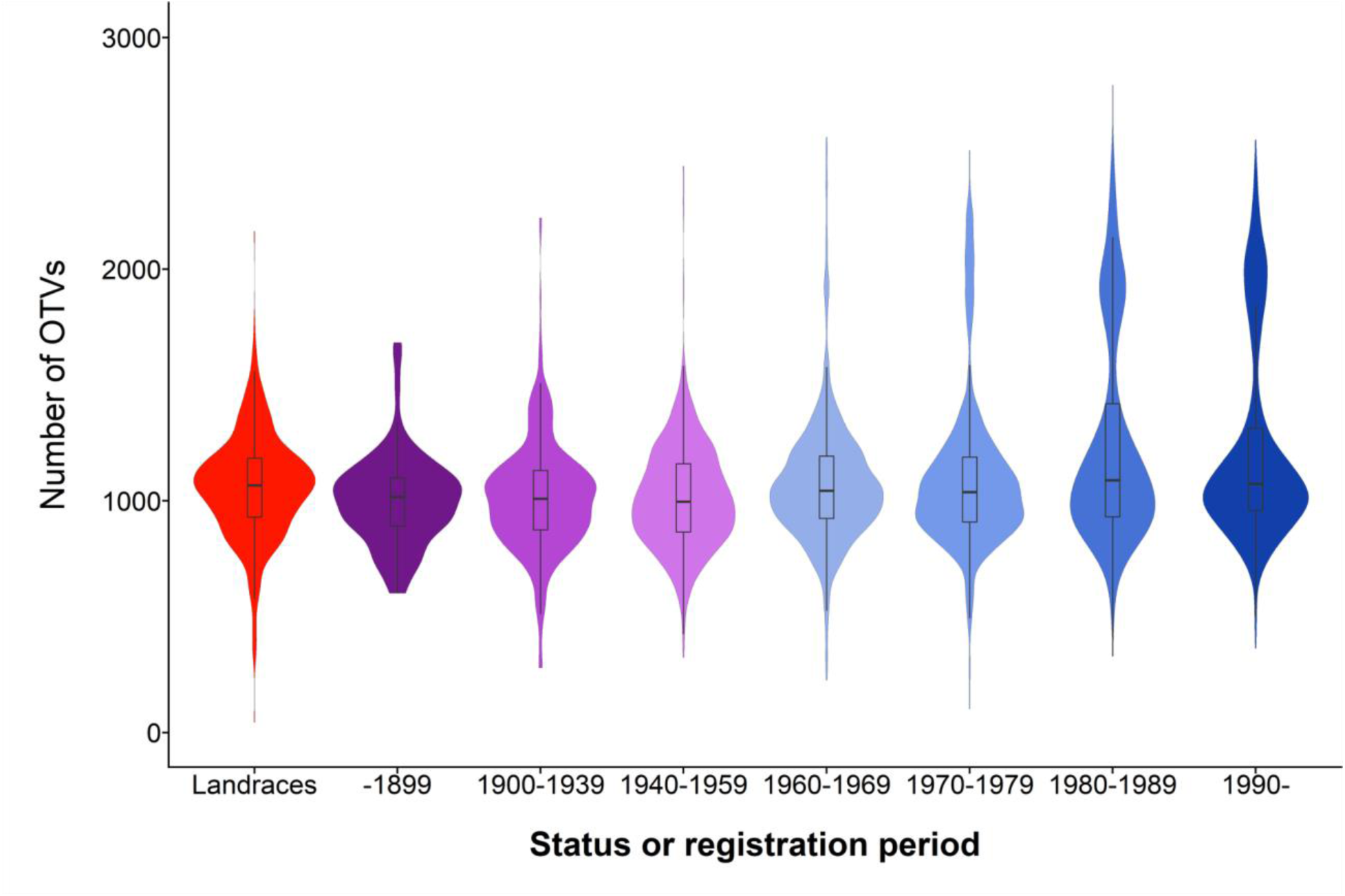
Frequency of occurrence of OTVs in wheat accessions according to the status or registration period. Landraces in red, traditional lines (before 1960) in purple, and modern varieties (from 1960 onward) in blue.

Altogether, these results brought new insights into the worldwide wheat genetic diversity diffusion and evolution. Recent selection and spread led to a modern germplasm that is highly unbalanced compared to the ancestral one found in landraces. Indeed, most of worldwide modern cultivars are related to the SEE, IBP and MED landraces, whereas NWE landraces contributed only to Western Europe modern germplasm and Asian landraces (SEA, CAA, INP and CAU) have been scarcely used in modern wheat breeding programs. In addition, the introduction of wild relatives in crop improvement has resulted in a significant amount of alien DNA within the genome of wheat varieties.

From a breeder’s perspective, the Asian germplasm represents a significant fraction of the worldwide diversity that remains largely unexploited. Nevertheless, as exemplified by the *Rht* dwarfing genes (23), as well as by disease resistance genes, such as *Fhb1* (24), these accessions are a valuable reservoir of novel alleles and genes. It is thus strategic to better characterize these genetic resources in order to exploit them efficiently into pre-breeding programs and fully benefit from their natural resistance to biotic and abiotic stresses.

It is now widely admitted that, because of intraspecific SVs, a single individual genome is not sufficient to describe accurately the whole species (25). Indeed, while some genes are shared between all individuals (the core genome), some others may be absent from one or more of them (the dispensable genome), the sum of all of the genes defining the pangenome (26, 27). First attempts to get access to the wheat pangenome have already been made (28). However, in the absence of a preliminary detailed phylogeographical study, the selected lines might not be representative of the worldwide diversity. We think that non-admixed landraces from each genetic groups described above can be considered as founder lines and would be the ideal candidates to characterize the wheat pangenome. After the completion of a high-quality reference sequence of the hexaploid wheat genome, this is the next challenge for the International Wheat Genome Sequencing Consortium (IWGSC).

## Acknowledgments

The authors would like to acknowledge the International Wheat Genome Sequencing Consortium and especially Rudi Appels (Murdoch University, Murdoch, Australia), Catherine Feuillet (Inari Agriculture, Boston, USA), Beat Keller (University of Zurich, Zurich, Switzerland), Sebastien Praud (Biogemma, Chappes, France), Ute Baumann (University of Adelaide, Adelaide, Australia), Hikmet Budak (Montana State University, Bozeman, USA), Jane Rogers (Eversole Associates, Bethesda, USA), and Kellye Eversole (Eversole Associates, Bethesda, USA). They are also grateful to the BreedWheat consortium leaders: Michaël Alaux (URGI, INRA, Versailles, France), Bernard Bejar (Céréales Vallée, Saint-Beauzire, France), Stéphane Lafarge (Biogemma, Chappes, France), Emmanuelle Lagendijk (INRA Transfert, Clermont-Ferrand, france), Jérémy Derory (Limagrain, Chappes, France), and Jacques Le Gouis (GDEC, INRA, Clermont-Ferrand, France). Many thanks to Marion Deloche and Lionel Bardy (GDEC, INRA, Clermont-Ferrand, France) for providing seeds, and to Michael Alaux and Thomas Letellier (URGI, INRA, Versailles, France) for hosting data. Axiom genotyping was conducted on the genotyping platform GENTYANE at INRA Clermont-Ferrand (gentyane.clermont.inra.fr). Sample seeds were provided by the Small Grain Cereals Biological Resources Centre at INRA Clermont-Ferrand (http://www6.clermont.inra.fr/umr1095_eng/Teams/Research/Biological-Resources-Centre).

## Funding

The research leading to these results have received funding from the French Government managed by the Research National Agency (ANR) under the Investment for the Future programme (BreedWheat project ANR-10-BTBR-03), from FranceAgriMer, French Funds to support Plant Breeding (FSOV), from Région Auvergne and from INRA. RDO was funded by a grant from the French Ministry for Research.

## Author contributions

Conceptualization: FB, SB, EP; Methodology: FB, SB, EP; Software: HR, FC; Validation: FB, SB, EP; Formal Analysis: FB, SB, RDO, SR, EP; Investigation: JK, HR, FC, EP; Resources: FB, IWGSC, BW, EP; Data Curation: HR, EP; Writing – Original Draft Preparation: FB, SB, EP; Writing – Reviewing & Editing: FB, SB, EP; Visualization: FB, SB, RDO, EP; Supervision: FB, EP; Project Administration: FB, EP; Funding Acquisition: FB, BW, EP.

## Competing interests

Authors declare no competing interests.

## Data and materials availability

Sample seeds of studied accessions can be requested under MTA at the INRA small grain cereals BRC or by using SIReGal website (https://urgi.versailles.inra.fr/siregal/siregal/grc.do).

## References and Notes

1. M. Heun et al., Site of einkorn wheat domestication identified by DNA fingerprinting. Science 278, 1312 (1997).

2. F. Salamini, H. Ozkan, A. Brandolini, R. Schafer-Pregl, W. Martin, Genetics and geography of wild cereal domestication in the Near East. Nat. Rev. Genet. 3, 429 (2002).

3. S. Lev-Yadun, A. Gopher, S. Abbo, The Cradle of Agriculture. Science 288, 1602 (2000).

4. S. Riehl, M. Zeidi, N. J. Conard, Emergence of Agriculture in the Foothills of the Zagros Mountains of Iran. Science 341, 65 (2013).

5. M. Feldman, in The World wheat book: A history of wheat breeding, A. P. Bonjean, W. J. Angus, Eds. (Lavoisier Publishing, Paris, 2001), pp. 3-56.

6. P. S. Baenziger, W. K. Russell, G. L. Graef, B. T. Campbell, Improving lives: 50 years of crop breeding, genetics, and cytology. Crop Sci. 46, 2230 (2006).

7. G. Conway, G. Toenniessen, Feeding the world in the twenty-first century. Nature 402, C55 (1999).

8. Food and Agriculture Organization of the United Nations, FAOSTAT Statistics Database. http://www.fao.org/faostat/en/#data/QC (2017)

9. F. Balfourier et al., A worldwide bread wheat core collection arrayed in a 384-well plate. Theor. Appl. Genet. 114, 1265 (2007).

10. A. Horvath et al., Analysis of diversity and linkage disequilibrium along chromosome 3B of bread wheat (Triticum aestivum L.). Theor. Appl. Genet. 119, 1523 (2009).

11. H. Rimbert et al., High throughput SNP discovery and genotyping in hexaploid wheat. PLoS ONE 13, e0186329 (2018).

12. International Wheat Genome Sequencing Consortium, Shifting the limits in wheat research and breeding through a fully annotated and anchored reference genome sequence. Submitted to Science, (2018).

13. J. Lachance, S. A. Tishkoff, SNP ascertainment bias in population genetic analyses: Why it is important, and how to correct it. BioEssays 35, 780 (2013).

14. S. Bouchet et al., Adaptation of Maize to Temperate Climates: Mid-Density Genome-Wide Association Genetics and Diversity Patterns Reveal Key Genomic Regions, with a Major Contribution of the Vgt2 (ZCN8) Locus. PLoS ONE 8, e71377 (2013).

15. Z. H. He, S. Rajaram, Z.Y. Xin, and G.Z. Huang, A History of Wheat Breeding in China. (CIMMYT, Mexico City, Mexico, 2001).

16. J. P. Didion et al., Discovery of novel variants in genotyping arrays improves genotype retention and reduces ascertainment bias. BMC Genomics 13, 1 (2012).

17. R. Hajjar, T. Hodgkin, The use of wild relatives in crop improvement: a survey of developments over the last 20 years. Euphytica 156, 1 (2007).

18. X. Cai, D. Liu, Identification of a 1B/1R wheat-rye chromosome translocation. Theor. Appl. Genet. 77, 81 (1989).

19. M. Helguera et al., PCR Assays for the Lr37-Yr17-Sr38 Cluster of Rust Resistance Genes and Their Use to Develop Isogenic Hard Red Spring Wheat Lines. Crop Sci. 43, 1839 (2003).

20. V. Kuraparthy et al., Characterization and mapping of cryptic alien introgression from Aegilops geniculata with new leaf rust and stripe rust resistance genes Lr57 and Yr40 in wheat. Theor. Appl. Genet. 114, 1379 (2007).

21. B. B. Wulff, M. J. Moscou, Strategies for transferring resistance into wheat: from wide crosses to GM cassettes. Front. Plant. Sci. 5, 692 (2014).

22. S. D. Tanksley, S. R. McCouch, Seed banks and molecular maps: unlocking genetic potential from the wild. Science 277, 1063 (1997).

23. P. Hedden, The genes of the Green Revolution. Trends Genet. 19, 5 (2003).

24. N. Rawat et al., Wheat Fhb1 encodes a chimeric lectin with agglutinin domains and a pore-forming toxin-like domain conferring resistance to Fusarium head blight. Nat. Genet. 48, 1576 (2016).

25. A. A. Golicz, J. Batley, D. Edwards, Towards plant pangenomics. Plant Biotechnol. J. 14, 1099 (2016).

26. H. Tettelin, D. Riley, C. Cattuto, D. Medini, Comparative genomics: the bacterial pan-genome. Curr. Opin. Microbiol. 11, 472 (2008).

27. R. K. Saxena, D. Edwards, R. K. Varshney, Structural variations in plant genomes. Brief. Funct. Genomics 13, 296 (2014).

28. J. D. Montenegro et al., The pangenome of hexaploid bread wheat. Plant J. 90, 1007 (2017).

29. J. C. Barrett, B. Fry, J. Maller, M. J. Daly, Haploview: analysis and visualization of LD and haplotype maps. Bioinformatics 21, 263 (2005).

30. S. Purcell et al., PLINK: A Tool Set for Whole-Genome Association and Population-Based Linkage Analyses. Am. J. Hum. Genet. 81, 559 (2007).

31. J. K. Pritchard, M. Stephens, P. Donnelly, Inference of population structure using multilocus genotype data. Genetics 155, 945 (2000).

32. D. Falush, M. Stephens, J. K. Pritchard, Inference of population structure using multilocus genotype data: linked loci and correlated allele frequencies. Genetics 164, 1567 (2003).

33. B. S. Weir, C. C. Cockerham, Estimating F-Statistics for the Analysis of Population Structure. Evolution 38, 1358 (1984).

34. J. Goudet, hierfstat, a package for r to compute and test hierarchical F‐statistics. Mol. Ecol. Notes 5, 184 (2005).

35. K. Jordan et al., A haplotype map of allohexaploid wheat reveals distinct patterns of selection on homoeologous genomes. Genome Biol. 16, 48 (2015).

36. S. Wang et al., Characterization of polyploid wheat genomic diversity using a high-density 90 000 single nucleotide polymorphism array. Plant Biotechnol. J. 12, 787 (2014).

37. S. Salvi, O. Porfiri, S. Ceccarelli, Nazareno Strampelli, the “Prophet” of the green revolution. J. Agric. Sci. 151, 1 (2013).

38. H. Zhonghu;, S. Rajaram, Z. Y. Xin, G. Z. Huang, A history of wheat breeding in China. (CIMMYT, Mexico, 2001).

39. C. Hao et al., The iSelect 9 K SNP analysis revealed polyploidization induced revolutionary changes and intense human selection causing strong haplotype blocks in wheat. Sci. Rep. 7, 41247 (2017).

40. A. Betts, P. W. Jia, J. Dodson, The origins of wheat in China and potential pathways for its introduction: A review. Quat. Int. 348, 158 (2014).

41. H. Tsujimoto, T. Yamada, T. Sasakuma, Pedigree of Common Wheat in East Asia Deduced from Distribution of the Gametocidal Inhibitor Gene (Igc1) and β-Amylase Isozymes. Jppn. J; Breed. 48, 287 (1998).

42. Y. Zhou et al., Uncovering the dispersion history, adaptive evolution and selection of wheat in China. Plant Biotechnol. J. 16, 280 (2018).

43. A. L. Olmstead, P. W. Rhode, Adapting North American wheat production to climatic challenges, 1839–2009. Proc. Natl. Acad. Sci. U.S.A. 108, 480 (2011).

44. Q. M. Paulsen, J. P. Shroyer, The early history of wheat improvement in the Great Plains. Agron. J. 100, S70 (2008).

45. S. Rajaram, M. Van Ginkel, in The World Wheat Book: A History of Wheat Breeding, A. P. Bonjean, W. J. Angus, Eds. (Lavoisier, Paris, France, 2001), pp. 579-610.

46. A. P. Bonjean, W. J. Angus, The World Wheat Book: A History of Wheat Breeding. (Intercept, 2001).

47. R. K. Bacon, in The World Wheat Book: A History of Wheat Breeding, A. P. Bonjean, W. J. Angus, Eds. (Lavoisier, Paris, France, 2001), pp. 469-478.

48. C. J. Peterson, R. E. Allan, C. J. Peterson, in The World Wheat Book: A History of Wheat Breeding, A. P. Bonjean, W. J. Angus, Eds. (Lavoisier, Paris, France, 2001), pp. 407-430.

49. B. F. Carver, A. R. Klatt, E. G. Krenzer, in The World Wheat Book: A History of Wheat Breeding, A. P. Bonjean, W. J. Angus, Eds. (Lavoisier, Paris, France, 2001), pp. 445-468.

50. R. H. Busch, T. Rauch, in The World Wheat Book: A History of Wheat Breeding, A. P. Bonjean, W. J. Angus, Eds. (Lavoisier, Paris, France, 2001), pp. 431-444.

51. R. Joukhadar, H. D. Daetwyler, U. K. Bansal, A. R. Gendall, M. J. Hayden, Genetic Diversity, Population Structure and Ancestral Origin of Australian Wheat. Front. Plant Sci. 8, 2115 (2017).

