## Supplementary material for "Worldwide phylogeography and history of wheat genetic diversity"

#### Materials and Methods

Wheat accessions

The worldwide diversity panel included 4,506 accessions originating from 105 different countries: 2,185 accessions from 33 different European countries, 143 from 28 different regions of the Russian Federation, 534 from 29 US states and 5 Canadian provinces, 946 from Asia, including 384 from 24 Chinese provinces, 316 from 14 South American countries, 207 from 20 African countries and 169 from Oceania. Pedigrees were known for 90% of the accessions. There were 59% of winter, 38% of spring and 3% of facultative types. Fourteen percent of these accessions (632) were landraces, 21% (965) corresponded to traditional cultivars (registered before 1960) and 51% were modern varieties (registered after 1960) (Data S1).

Genotyping

The 4,506 accessions were genotyped using a high-density Affymetrix Axiom SNP array containing 280,226 genic and intergenic SNPs (*11*). Genotyping was conducted on the Affymetrix GeneTitan system according to the procedure described by Affymetrix (Axiom® 2.0 Assay Manual Workflow User Guide Rev3). Allele calling was carried out using a modified version of the Affymetrix proprietary software packages Affymetrix Power Tools (APT) and SNPolisher™ (http://www.affymetrix.com/estore/partners_programs/programs/developer/tools/devnettools.affx) to take into account the specificities of the wheat genome. For all SNPs, HomRO and HomFLD were calculated (http://media.affymetrix.com/support/developer/downloads/Tools/SNPolisher_User_Guide.pdf).

The HomFLD filter was set to 3.6. As a first step, all the probesets were processed with a mild inbred penalty equal to 4 on all the samples. As a second step, the SNPs failing the QC criteria ("Other" and "NoMinorHom") were reprocessed using an inbred penalty of 16. Probesets classified as OTVs by SNPolisher were analyzed with OTV_caller in the two steps.

Haplotyping

Missing data were estimated using the Beagle software. Lines with more than 5% missing data were discarded from this analysis. Haplotypic blocks and specific haplotypes contained in these blocks were defined in 4,403 lines using Haploview algorithm (*29*) implemented in Plink software (*30*). In each haplotypic block, 95% of pairwise measures of marker LD (D') were superior to 0.98.

Structure

In order to define a relevant number of genetic groups in 632 worldwide bread wheat landraces and to assign lines to genetic groups, we used haplotypes defined above and the algorithm implemented in STRUCTURE software (*31*). We assumed a single domestication event and restricted our analysis to the correlated frequency model (*32*). We set other parameters at their default values using the admixture model and infer ALPHA option. We used a 10,000 burn-in period and 10,000 iterations. Allele frequencies in each of the K clusters (from 2 to 15) were estimated, and the percentage of genome derived from each cluster was estimated for each accession. Pairwise differentiation indexes Fst (*33*) were calculated with r-Hierfstat (*34*) to estimate the distances between groups.

#### Supplementary Text

Genotyping

SNPs were classified in six main categories according to cluster patterns produced by the Affymetrix software: Polymorphic High Resolution (PHR) Off-Target Variants (OTV), Monomorphic High (MHR), No Minor Homozygous (NMH), Call Rate Below Threshold (CRBT) and Others. A set of 113,457 high quality SNPs showing less than 2% missing data was selected (Data S2). This comprised 99,333 PHR bi-allelic SNPS as well as 14,124 OTVs, *i.e.* markers that detect both nucleotide polymorphisms and presence-absence variations. The genomic position of these SNPs was determined using the IWGSC RefSeq v1.0 (*12*). The distribution of markers on the homoeologous genomes was consistent with previous studies (40% on the A-, 48% on the B- and 12% on the D-genomes) (*35, 36*).

Haplotypes

We parsed the bread wheat genome in 8,741 sizable regions over which there was little evidence for historical recombination and within which only a few common haplotypes were observed (Data S3). The number of haplotype blocks was highly correlated with the overall number of SNPs per chromosome (Pearson correlation coefficient R=0.79, p=2E-5). The average number of haplotypes per block was 4, ranging from 2 to 20.

The median size of haplotype blocks is 105 kb, with 85% of the blocks being shorter than 1 Mb. The mean size is 863 ± 4,595 kb. This huge standard deviation reflects the structural partitioning of wheat chromosomes. Indeed, the size was found to be different according to the five chromosomal regions defined by IWGSC, as expected from the overall recombination pattern observed in wheat (*12*). In telomeric highly-recombinogenic R1 and R3 regions, the mean size was 226 ± 47 kb and 338 ± 86 kb, respectively; in the R2a/b regions, 2,239 ± 87 kb; and in the C region, 15,117 ± 3,387 kb (Figure S1).

Ascertainment bias

Mean similarities between accessions were 0.69 (sd = 0.07), 0.70 (sd = 0.05) and 0.49 (sd = 0.07) using 113,457 SNP, 58,602 pruned SNPs or 8,741 haplotypes respectively. Similarities calculated with haplotypes were much lower. Correlation was higher between similarities calculated with 113K or 59K pruned SNP ($R^2 = 0.95$), 8K haplotypes or 59K pruned SNP ($R^2 = 0.96$) than between similarities calculated with 113K SNP or 8K haplotypes ($R^2 = 0.88$) (Figure S2). Similarity differences when calculated with SNP or haplotypes were very low for individuals that were very similar, intermediate when individuals were very different and high when individuals were moderately different (Figure S3). Haplotypes may have more power to discriminate individuals thanks to the high number of alleles.

Population structure

In order to define a relevant number of genetic groups in 632 worldwide bread wheat landraces and to assign each line to each genetic group, haplotypes were used to analyze the population structure for the 632 landraces with the STRUCTURE software.

We first ordered groups of each run so that they have the maximum number of individuals in common. Then we calculated assignment discrepancy as the proportion of individuals with assignment >0.8 that are assigned to different groups between runs. We observed that for a number of groups ranging from 6 to 9, assignments were quite stable (Figure S4).

To avoid the bias due to the difficulty to find correspondence between small groups obtained in different runs, we also looked at the correlation between similarities calculated using individuals allele frequencies estimated by STRUCTURE, *i.e.* the cross product of group estimated allele frequencies and assignment to those groups (Figure S5). Those are quite stable between runs until K=8.

We chose to describe the panel using eight groups as it discriminates the Iberian Peninsula from Mediterranean lines and Caucasian lines from the Western Asia, which sounded biologically and historically relevant to notice (Figure S6). At K=2, wheat landraces were separated according to their European *vs.* Asian origins. At K=3, the Asian cluster was divided in two clusters: South East Asian (SEA) and Central Asia-Africa (CAA). At K=4, European landraces were split between the North West (NWE) and the South-East Europe (SEE) clusters. At K=5, the Mediterranean Basin cluster (MED) was separated from North West Europe. At K=6, the Indian Peninsula cluster (INP) was derived from Central Asian cluster. At K=7, the Iberian Peninsula cluster (IBP) separated from MED. At K=8, the Caucasus cluster (CAU) split from the CAA cluster.

The geographical origin of landraces from these eight sub-populations is given in Data S1 and the composition of the eight sub-populations in Table S1. CAA landraces were from Western Asia (30%) (mostly Oman, Saudi-Arabia), Southern Asia (22%) (Pakistan, Iran) and Central Asia (9%) (Tajikistan), Northern Africa (20%) (Sudan) and Western Africa (11%) (Mali, Niger). They were mostly (86%) spring type. CAU landraces were from Western Asia (68%) (Azerbaijan, Turkey, Georgia, Armenia) and Eastern Europe (22%) (Russia). Most INP landraces were from Southern Asia (83%) (India, Pakistan, Nepal). They were exclusively spring type. Most SEA landraces were from Eastern Asia (98%), 85 % from China and 10 % from Japan. NWE landraces came from Western Europe (60%) (France, Germany), Northern Europe (15%) (United Kingdom) and Southern Europe (12%) (Spain). They were mostly (84%) winter type. MED landraces came from Western Asia (29%) (Syria, Turkey), Southern Europe (27%) (Spain, Italy), Northern Africa (17%) (Morocco, Tunisia) and Western Europe (10%) (France). They were mostly (85%) spring type. IBP landraces were from Spain (75%) and Portugal (25%). They are mostly (75%) spring type. SEE landraces were from Eastern Europe (35%) (Russia, Ukraine, Hungary), Western Europe (14%) (France), Western Asia (13%) (Turkey, Azerbaijan) and Northern Europe (11%) (UK, Sweden, Finland).

While showing the strong geographical structure existing within wheat landraces, the projection on the two first axes of a Principal Component Analysis based on the Sokal and Michener dissimilarity matrix calculated with 8,741 haplotypes revealed a continuum of genetic diversity between populations, consistent with the high admixture rate (Figure S7).

SEE, NWE and MED were the most admixed (mean assignment rate= 0.71, 0.68 and 0.66, respectively) and the most diverse groups (He= 0.44, 0.45 and 0.47, respectively) (Table S2). INP, CAU, SEA, and IBP were the least admixed (mean assignment rate= 0.77, 0.80, 0.88 and 0.89, respectively) and the least diverse groups (He= 0.35, 0.40, 0.34 and 0.30, respectively). SEA and IBP were the most differentiated groups (Fst = 0.32 and 0.36 respectively compared to 0.2 on average for the other groups). SEE, NWE and MED were not highly differentiated from each other (pairwise Fst = 0.10 on average). Global Fst was 0.4.

Spatial structuration of the worldwide diversity

After haplotypic blocks inference and filtering for lines with more that 5% missing data, we analyzed a worldwide diversity panel of 4,403 accessions.

The phylogenetic analysis defined eleven groups within the worldwide diversity (Figure 1, Data S1 and Table S3). Group I was composed of Ukrainian, Hungarian, Russian, Yugoslavian and Romanian modern winter type (93%) lines. Group II was composed of French, Canadian, German, North American and Swedish, accessions that are mostly (80%) spring type. Group III was composed of North American, Russian, Hungarian, Ukrainian and Romanian, mostly winter type (86%) accessions. Group IV was composed of New World (America, Australia), French and Spanish accessions. Group V was composed of Chinese, Italian, Yugoslavian and Hungarian modern lines. Group VI was composed of CIMMYT, New World and Chinese modern spring type (93%) lines. Group VII was composed of Canadian, Chinese and American modern exclusively spring type lines. Group VIII was composed of Asian, mostly (77%) spring accessions, including most INP, CAA and CAU landraces. Group IX was composed of Chinese and Japanese accessions, including most SEA landraces. Group X was composed of French and British modern winter type (90%) lines. Group XI was composed of French and German modern winter type (96%) lines.

Interestingly, all non-admixed (assignment > 0.6) CAA, INP and most CAU (92%) landraces were found in group VIII, and most of the non-admixed SEA landraces (98%) in group IX. Non-admixed NWE landraces were distributed in groups XI (56%), X (24%), V (10%) and IV (9%). All non-admixed MED and IBP landraces were in group IV. Non-admixed SEE landraces clustered in groups III (52%), II (19%) and IV (18%). Note that no non-admixed landraces were found in group VI and VII and only a few (8) in group I.

The most differentiated group was VII (Fst > 0.3 with all other groups). VII modern varieties were even more differentiated than VII traditional cultivars (Table S5). IX traditional cultivars were highly differentiated to all groups except group VIII (Fst = 0.11). Note that IX modern varieties were less differentiated than IX traditional cultivars. They were probably admixed between different ancestral groups. This suggests that IX traditional cultivars did not participate to any elite lines except IX elites. VIII traditional and VIII elites were very close to each other (Fst = 0.02). VIII traditional were less differentiated to worldwide elite germplasm than IX traditional. They probably slightly contributed to modern elite germplasm although IX traditional did not.

A detailed analysis was conducted for different geographical regions and countries to determine the proportion of sampled accessions according to their clustering in eleven genetic groups, as indicated on figure 2 (Figure S8 to S16).

*Europe*

Accessions from European countries were mainly found within five different groups, namely groups I, II, III, X and XI (Figure S8). North Western European accessions were clustered in groups X and XI, while accessions from Eastern European countries were mainly found in groups I and III. This strong structure between North Western and South Eastern European wheat genepools may be related to the historical roads used by the first farmers during the spread of agriculture. In particular, the Danubian road can be a hypothesis to explain this geographical separation between groups X and XI on the one hand, and groups I and III on the other hand. The split of Western Europe accessions between groups X (France, UK, Belgium…) and XI (Germany, The Netherlands, Poland, Denmark...) may be related to old selection footprints for specific climatic conditions such as oceanic/continental climates. Finally, some accessions clustering in group II (pink) can be found in the northern part of Europe (Scandinavia). This group contained mainly spring type accessions well adapted to specific day length and vernalization requirements in Northern European countries (Norway, Sweden, Finland, and Ireland).

*Mediterranean Basin*

The material from the Mediterranean Basin differed from the European material (Figure S9). Indeed, four major groups (IV, V, VI, and VIII) were observed. Group I was also represented but only in Balkans area (former Yugoslavia, Macedonia, Bulgaria). The level of diversity and admixture was high in this region at the cross of different migration roads. The Mediterranean road from Turkey, Greece across Italy, France and Spain was likely related to group IV while the North African road might have given raise to groups VI and VIII. Finally, we can notice a small group V with few accessions, highly specific from the Adriatic sea (Italy, Croatia, Former Yugoslavia, Albania, Greece, ..) that might reflect high selection pressures to very specific environmental conditions such as drought resistance.

*Middle East & Central Asia*

This large Middle East & Central Asia geographical area corresponds to the Cradle of Wheat and the starting point of the main historical migration roads. It includes the center of diversification of wheat genepool. We observed accessions from groups I, III, IV, VI, and VIII (Figure S10). Groups I and III seems to be originating from the region of the Caspian Sea, with Caucasus (Armenia, Georgia, Azerbaijan, Russian Federation) on the West and former Persia

(Kazakhstan, Turkmenistan, Uzbekistan, Tajikistan) on the East. It is worth noting that group I was also well represented on the other side of the Black Sea, as previously observed on Figure S8. Accessions from group VIII mainly corresponded to landraces from the CAA and CAU populations. Finally, a majority of accessions belonging to groups IV and VI, previously observed in figure S9, were from Turkey, Iraq, Iran, Israel, Lebanon, and Jordan, that is the former Fertile Crescent Area.

*South-Eastern Asia*

South-Eastern Asia diversity was largely dominated by five main groups (Figure S11). Group VIII was the most prevalent one in the Indian Peninsula (Pakistan, India, Nepal, Bhutan), as well as in Myanmar, Philippines, and Indonesia and comprised mainly landraces from the INP population. In contrast, China, Japan and Korea presented a high proportion of accessions from group IX that corresponded to SEA landraces, while one third of recent Chinese cultivars clustered in group V. The distribution of accessions between groups VI, VIII and IX (the first two being also observed at high frequency in Middle East and Central Asia) varied along the Southern extremity of the Silk Road used by first farmers from the Fertile Crescent through Central Asia, Afghanistan and Xinjiang (North-West China). The importance of group V in Chinese modern cultivars and in Italy (Figure S9) can be explained by intensive uses of Italian germplasm in Chinese breeding programs during the second part of 20$^{th}$ century (*37-39*). Indeed, several famous Italian accessions, such as Mentana, Mara, Funo, Libellula, or Virgilio that were developed by the Italian wheat breeder Strampelli and used as parent to create a large number of Chinese cultivars during the 1960s and the 1970s belong to group V. Finally, group III observed in Caucasus was also found in Mongolia, Republic of Korea, Japan, China, and Russia. The occurrence of this group supports the hypothesis of a secondary Northern way of introduction from Eurasia through Southern Siberia and Mongolia, along the Eurasian Steppe road (*40*, *41*).

Figure S12 presents a more precise description of wheat diversity in China, based on a set of 384 accessions originating from 24 different Chinese provinces. As previously observed on Figure S11, these accessions were clustered in five main groups. The frequency of group VIII was higher in western provinces, such as Xinjiang and Tibet, closed to the Indian Peninsula where this group VIII was dominant. It only included landraces. Accessions clustered in group V were mainly observed in central part of China (Sichuan, Shaanxi, Henan, Hubei, Anhui,..), while group IX was dominant in North-East China (Shanxi, Hebei, Beijing, Tianjin, Shandong), together with accessions of group I. Finally, accessions of group VI were mainly observed in Northern provinces (Gansu, Ningxia, Liaoning). When considering the four main wheat growing zones in China, we observed a quite good correlation between these regions and the different groups. Indeed, autumn-planted spring wheat region was related to group IV while group IX was much more connected to winter habit and facultative wheats regions. Similarly, group VIII was mainly observed in spring-planted spring wheat and autumn-planted winter wheat area (Western part of China) while group VI was specific to spring-planted spring wheat region. These results reflect the effects of environmental selection pressures on haplotypic diversity and are consistent with those described by Zou and collaborators (*42*).

*Africa*

Despite the fact that not all African countries were represented in our panel, the African continent was largely dominated by the three main groups observed in Mediterranean Basin (IV and VI) and Middle East (VIII) (Figure S13). This diversity may have been shaped along ancient human migration roads from the Fertile Crescent (along the Nile, Tigris and Euphrates rivers, from the Red Sea to the Persian Gulf : Iraq, Syria, Lebanon, Cyprus, Jordan, Israel, Palestine, Egypt, the southeastern fringe of Turkey and the western fringes of Iran) to Northern Africa (Egypt, Tunisia, Algeria, Morocco) (groups IV and VIII). Group VI contains mainly (93%) modern lines. It may be recent introductions of ICARDA and CIMMYT wheat breeding materials to Eastern (Ethiopia, Kenya) and Southern (Zimbabwe, South Africa) parts of the continent.

*America*

Structure of wheat genepool diversity in the American continent is presented on figure S14. At the whole continent level, accessions were clustered in five main groups. Group IV mainly observed on the Eastern coasts of both North and South America (USA, Venezuela, Brazil, Uruguay, and Argentina) is likely related to the first introduction of wheat landraces by Spaniards from Iberian Peninsula to Eastern American coast during the 16$^{th}$ century. The presence of group III can be explained by the introduction of wheat accessions related to Eastern Europe lines and more adapted to northern environmental conditions and uses by European migrants such as the Mennonites who brought the Turkey Red Wheat to Kansas in the late 1800's during the settlement of the United States in North-America and Canadian provinces (*43, 44*). The introduction of spring wheat from Scandinavia and Northern part of Europe to Canada can explain the importance of group II in this country. Group VI differentiation reflects the remarkable breeding program developped at CIMMYT in Mexico since 1940 (*45*). This International organism together with ICARDA in Alep (Syria) produced and spread new cultivars in developing countries. They played a key role during the Green Revolution and the generalization of semi-dwarf high yielding varieties. Our results clearly indicates that CIMMYT and ICARDA mostly used Mediterranean accessions (groups IV and VI), and to a lower extent Middle East accessions of group VIII, to create then release new lines all over the world, to America, Oceania and Africa in particular. The very small group VII mostly observed in Canada and some Northern states of USA could be derived from recent Canadian breeding program using CIMMYT material.

A more precise description of wheat diversity in the USA, based on a set of 359 accessions originating from 30 different states, is presented on figure S15. As previously observed on figure S14, these accessions were clustered in five main groups that show a strong geographical structuration and are mostly related to the four main wheat types grown in this country, namely Soft White, Soft Red Winter, Hard Red Winter and Hard Red Spring wheats (*46*). Group IV was dominant in the Eastern part of the country (Missouri, Indiana, Michigan, Ohio, New-York, Georgia, North and South Carolina). This area is overlapping with the region corresponding to the US soft winter wheat pool (*47*). High frequencies of this group IV was also observed in Washington, Oregon and Idaho, where the soft white wheats are produced (*48*). By contrast, the highest frequencies for group III were observed in central and western states (Texas, Oklahoma, Kansas, Nebraska, Colorado, Utah) where the hard winter wheats are grown (*49*). Groups I and VII, which are dominant in Canada, showed the highest frequencies in Minnesota, North and South Dakota, Montana, near of Canadian border. These two groups can be related to the US hard red spring wheat pool (*50*). Finally, accessions gathered in group VI which can be observed in western part of the USA, are generally more recent cultivars with CIMMYT lines in their pedigrees. The first introductions of wheat in the USA may have occurred on the eastern coast at the beginning of the 17$^{th}$ century. A more important introduction of Mediterranean material occurred during the 19th with accession such as Mediterranean, Purple Straw or Red May. At the end of the 19th century, they appeared in the pedigree of better-adapted lines such as Goldcoin, Harvest Queen, Thorne, Chancellor, Atlas, Knock, Benhur, Arthur or Caldwell. All these accessions belong to group IV. During the progressive settlement in Western states of the country, soft wheat accessions from group III appeared to be non-adapted to drought or winter stress. So, new accessions from Eastern European pool, such as Turkey Red or Crimean from Ukraine, were introduced during the second part of the 19th century to create new cultivars such as Kharkov, Malakof, and varieties with higher yield potential such as Cheyenne, Nebred, Comanches, Wichita, Triumph, Centurk, Newton or Tam. All these accessions belong to group III. Soft winter wheats from Europe appeared not adapted to Northern states conditions. So, during the 19th century, settlers used spring types previously introduced from Scandinavia to Canada, such as Red Fife, Preston or Marquis to develop new cultivars as for instance Lee or Ceres, which all clustered in group II. Later, in the second part of the 1900s, introgressions of CIMMYT material in modern breeding programs produced more productive spring cultivars such as Justin, Chris, Era, Olaf or Len, all belonging to group VI. Some spring type European accessions from group III were at the origin of famous soft white wheat in Pacific Northwest region (Washington, Oregon, Idaho), such as Little Club introduced from California, Baart or Defiance. After the Second World War, introduction of CIMMYT wheat lines in plant breeding programs may also explain the importance of group VI in this Western part of the USA. The distribution of wheat diversity in North America corresponds to progressive introductions of specific Mediterranean and European materials and now reflects different cultivation systems and climates.

*Oceania*

Wheat accessions from Oceania were clustered in four main groups: groups II and X from Europe and groups IV and VI from the Mediterranean area and widely used by CIMMYT in recent breeding programs (Figure S16). This clustering is consistent with the evolution of Australian wheat as described by Joukhadar and collaborators (*51*). Indeed, wheat was first introduced in Oceania by the First Fleet in 1788. During the 1800s, growers used some famous lines from Europe and Mediterranean Basin, such as Fife (group II) or Purple Straw (group IV), then from Africa, like Gluyas (group IV). These lines were intensively used by breeders in the first half of the 20$^{th}$ century to develop the first commercial Australian varieties such as Federation, Bencubbin, Insignia, Olympic, Halbered or Spears which are all clustered in group IV. Since 1970, Australian wheat breeding has heavily relied on CIMMYT material to develop new famous cultivars such as Gabo, Gamenya, Condor, Egret, and Janz, which are all clustered in group VI. Therefore, groups IV and VI can be related to the main periods defined by Joukhadar et al. (*51*): group IV from the second period (1921-1970) and group VI from the third period (1971 onward). Interestingly, it seems that these groups also define the two main Australian wheat classes, namely white wheat (group IV) and hard wheat (group VI).

Evolution of the worldwide diversity through time

The level of diversity calculated with haplotypes through time is similar in landraces, traditional accessions and modern lines (0.52, 0.50, and 0.51, respectively), globally in the collection and in most groups (Figure S17). Although 9% of alleles were lost in modern varieties compared to landraces, 8% of new alleles were observed in modern lines that were enriched in structural variations. Although 7% of alleles that were specific to landraces contained OTVs, 19% of the alleles that were specific to modern material contained OTVs.

The proportion of the eleven groups appeared to be different according to the status of accessions: landraces, traditional cultivars (registered before 1960), and modern varieties (registered after 1960) (Figure 2B). Indeed, while groups IV, VIII and IX were dominant in landraces (28, 28 and 14%, respectively), their proportion decreased in traditional and modern lines, group VIII being almost absent for the latter. Concomitantly, I, VI, X and XI that were minority groups in landraces became dominant in modern varieties (15, 16, 17, and 12%, respectively). This can be explained by important breeding programs in Eastern Europe (group I), Western Europe (groups X and XI) as well as CIMMYT and ICARDA (group VI) since the 1960s.

To go a little bit further in the analysis of the shift that occurred in the genetic origin of wheat genepool through time, we studied these data in different geographic regions (Figure S17).

In Eastern Europe, landraces and traditional accessions that cluster mainly with group III (58 and 64%, respectively) were replaced by modern lines that mainly cluster with group I (60%). In Southern and Western Europe, group IV was dominant in landraces (68 and 52%, respectively) but was replaced in traditional cultivars and modern varieties by group V in Southern Europe (55 and 40%) and groups X and XI in Western Europe (33 to 48%). In Northern Europe, we observed a gradual shift from groups XI and III in landraces (43 and 33%, respectively), to groups XI (50%) and II (27%) in traditional, to a dominant group X (55%) in modern lines.

In Eastern Asia, while landraces, traditional and modern accessions mainly cluster with group IX (36 to 75%), we observed an increased proportion of the other groups, especially group

IV, absent in landraces and present in traditional and modern lines (16 and 14%, respectively). In Central Asia, group VIII that was dominant in landraces (63%) decreased in traditional (43%) to disappear almost completely from modern accessions (5%). Concomitantly, we observed an increase of group III in traditional (29%) and group I in modern lines (62%). In Southern Asia, the landrace- and traditional-dominant group VIII (88-90%) was replaced by group VI in modern varieties (68%). In Western Asia, although landraces mainly clustered with group VIII (45%) and IV (34%), modern lines mainly cluster with group VI (49%).

In Central-Eastern China (winter habit and facultative wheat,) although group IX was the main group in landraces (97%), traditional (91%) and modern (52%) accessions, we observed an increase of group I in the most recent varieties (29%). In South East China (autumn-planted spring wheat), group V gradually replaced the landrace-dominant group IX (81%) in traditional and modern lines (33 and 65%, respectively). Our panel contained only landraces from Western China (spring-planted spring and autumn-planted winter wheat). In North-Eastern and Central China (spring-planted spring wheat), we observed a complete change in the main groups, moving from III and IX in landraces (33 and 50%, respectively) to V and VI (50 and 20%, respectively) in modern varieties.

In Africa, although landraces mainly cluster with groups IV and VIII (45 and 39%, respectively), modern lines mainly cluster with group VI (72%).

In the New world, no accessions were considered as landraces. We thus only compared traditional cultivars and modern varieties, when possible.

In Canada, traditional cultivars mainly clustered with group II (47%) while groups VII (41%) and VI (28%) were dominant in modern varieties. In Northern USA (Hard Red Spring wheat), we observed an increase in group VI, from 14% in traditional to 56% in modern lines. A similar trend was found with group III in Mid-Western states (Hard Red Winter wheat), from 33% in traditional to 63% in modern. Interestingly, no significant change was observed in Southern and Eastern states (Soft Red Winter wheat) and North-Western states (Soft white wheat), the group III and IV dominating the former, and the group IV being the main group in the latter.

In Central America and on the Western Coast of South America, the absence of traditional cultivars in our panel did not allow the comparison. But on the Eastern Coast of South America, while traditional and modern lines mainly cluster with group IV, a noticeable increase in the proportion of group VI was observed (from 11 to 25%).

In New Zealand and Western Australia, the panel did not contain traditional cultivars. In Eastern Australia, group IV present in traditional (33%) disappeared almost completely from modern and was replaced by group VI (82%). A similar trend was observed in Southern Australia where group VI that was absent in traditional appeared in modern varieties (40%).

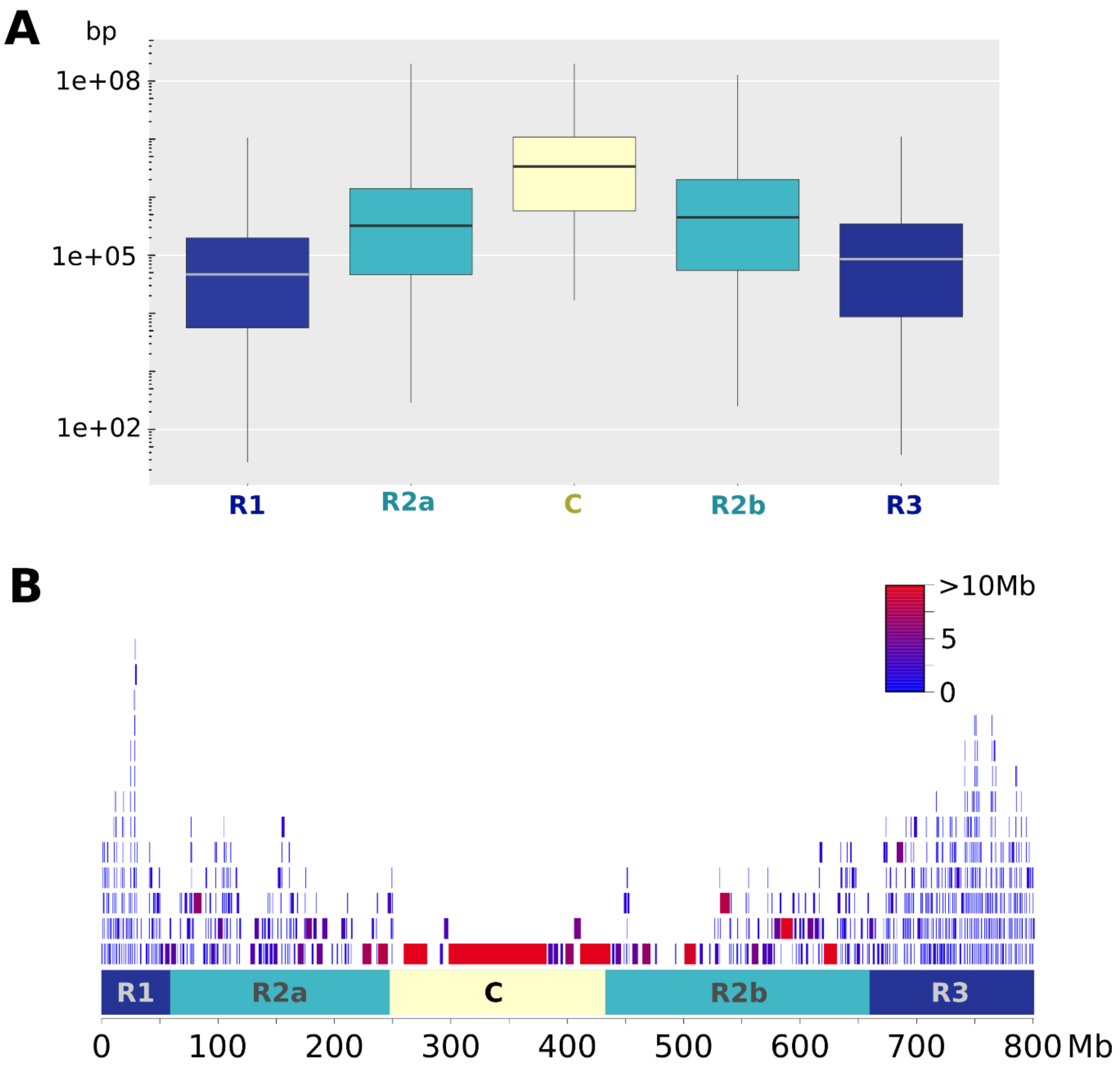


**Fig. S1. Size difference of haplotypic blocks along wheat chromosomes.**

(A) Boxplot of sizes in the R1, R2a, C, R2b and R3 regions of wheat chromosomes. (B) Distribution of haplotypic blocks along chromosome 2B. The block color corresponds to its size.

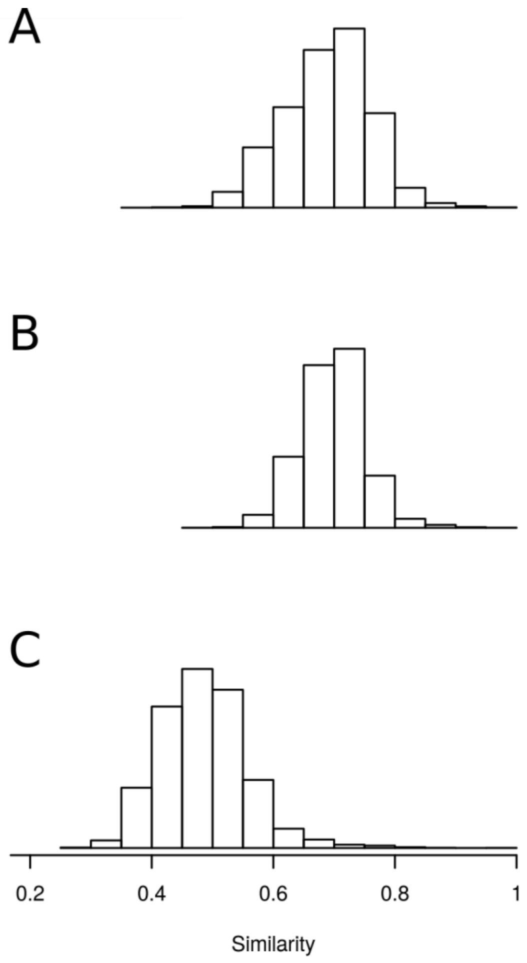


**Fig. S2. Distribution of similarities between 632 landraces calculated with different marker sets.**

(A) 113,457 SNPs; (B) 58,602 pruned SNPs; (C) 8,741 haplotypes.

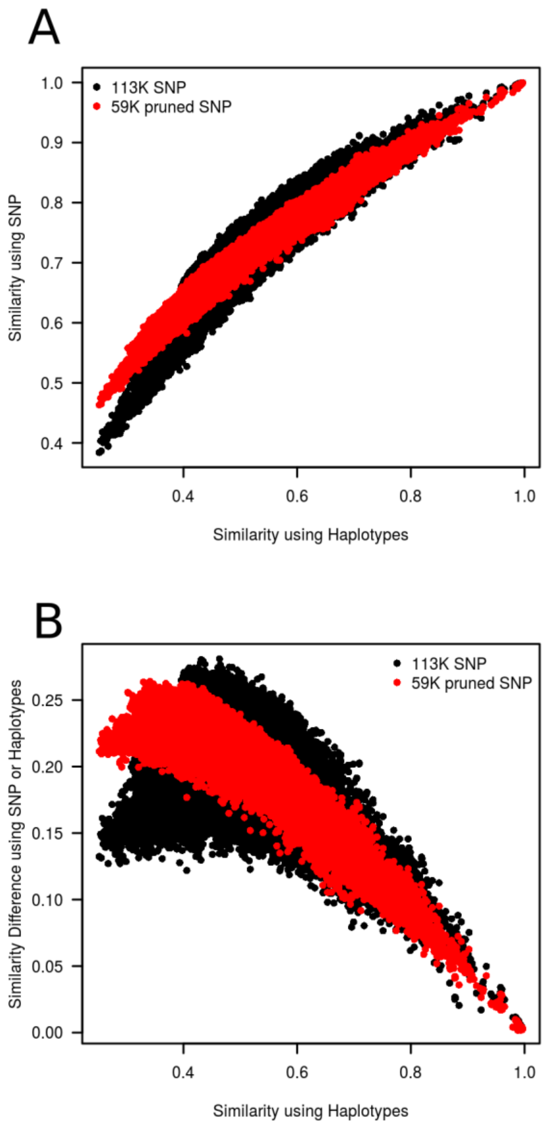


**Fig. S3. Comparison of similarities estimated with SNPs and haplotypes.**

**(**A) Correlation of similarities calculated with different marker sets: 8,741 haplotypes vs. 113,457 SNPs in black; 8,741 haplotypes vs. 58,602 pruned SNPs in red.

(B) Difference of similarity calculated with different marker sets as a function of similarity calculated using haplotypes: 8,741 haplotypes vs. 113,457 SNPs in black; 8,741 haplotypes vs. 58,602 pruned SNPs in red.

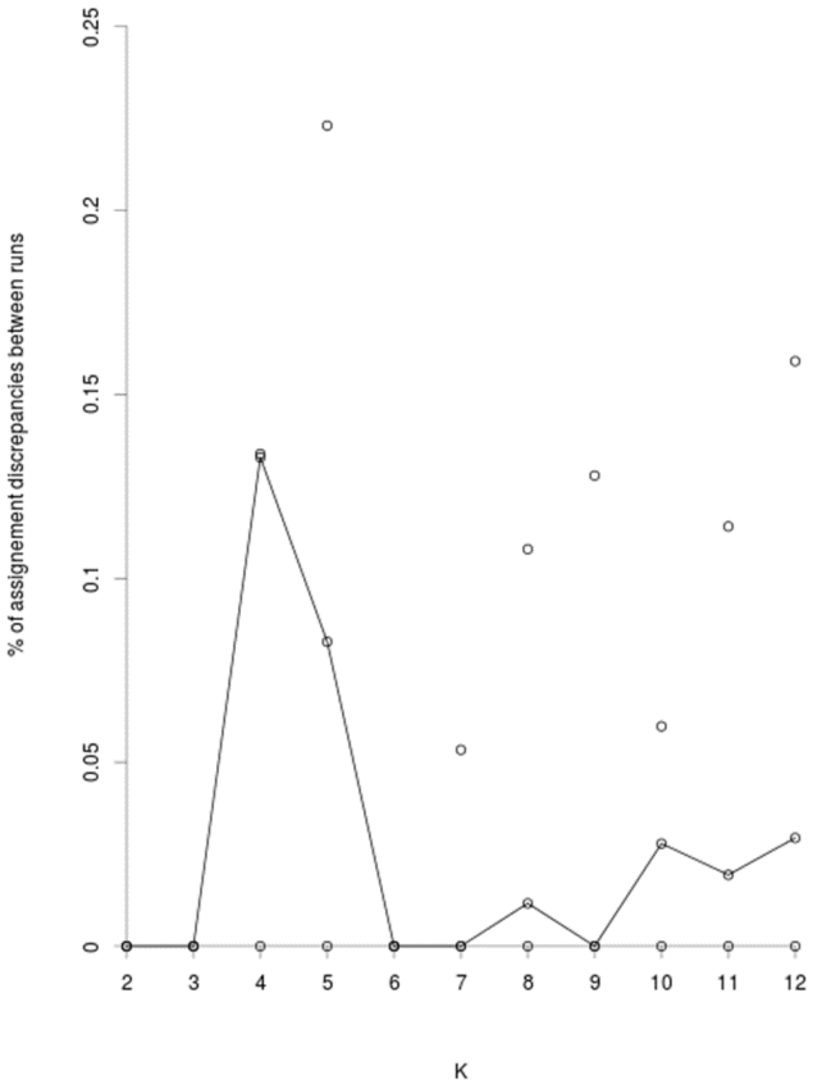


**Fig. S4. Stability of dissimilarity matrixes between the 632 landraces in different STRUCTURE runs.**

Similarities were calculated using group allele frequencies and assignment of individuals to groups obtained with STRUCTURE software. The number of groups ranges from 2 to 12. Five runs were implemented.

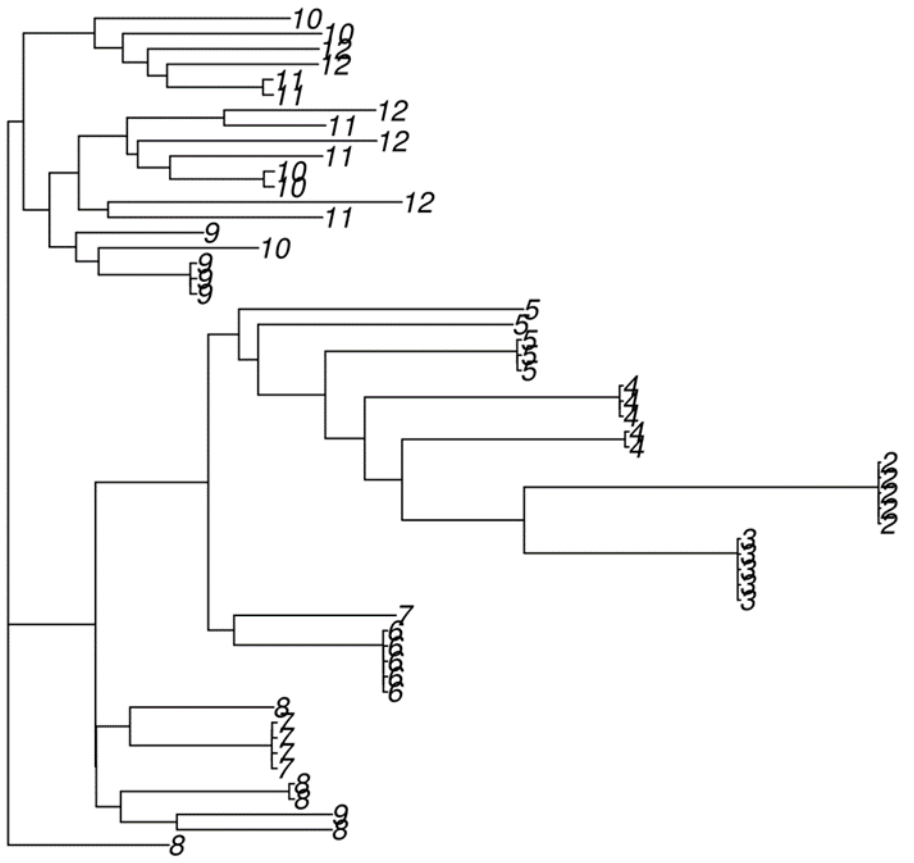

**Fig. S5. Stability of dissimilarity matrixes between the 632 landraces in different STRUCTURE runs.**

Five runs, and K ranging from 2 to 12. Dissimilarity matrixes were calculated using group allele frequencies and assignment of individuals to groups obtained with STRUCTURE software.

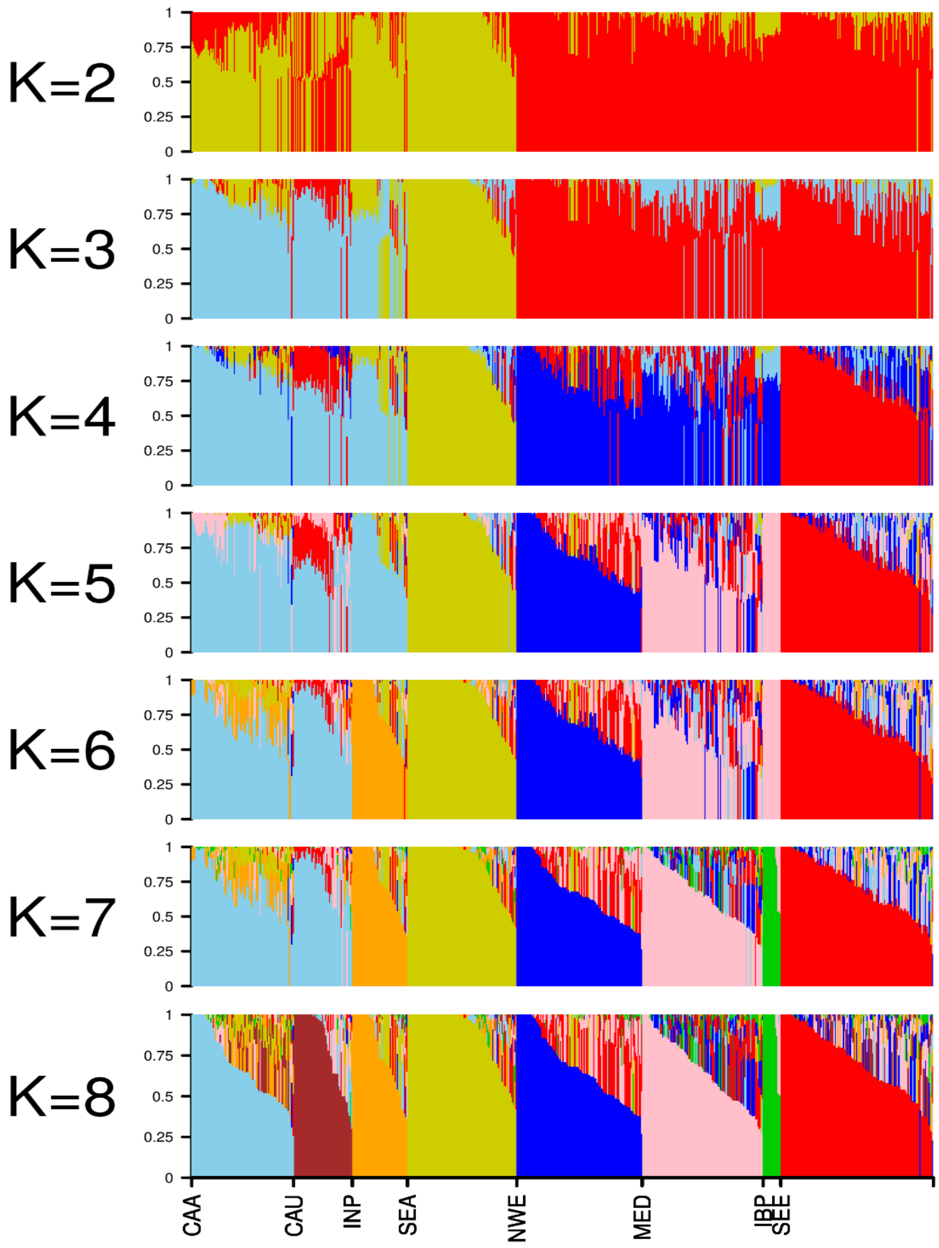


**Fig. S6. Assignment of 632 landraces to genetic groups.**

Structure was run using 8741 haplotypes. The number of genetic groups ranged from K = 2 to 12. Five runs were implemented for each K value. Assignments obtained by the run with the highest likelihood is represented.

Cyan: Central Asia & Africa (CAA); Brown: Caucasus (CAU); Orange: Indian Peninsula (INP); Yellow: South-East Asia (SEA); Blue: Nort West Europe (NWE); Pink: Mediterranean basin (MED); Green: Iberian Peninsula (IBP); Red: South East Europe (SEE).

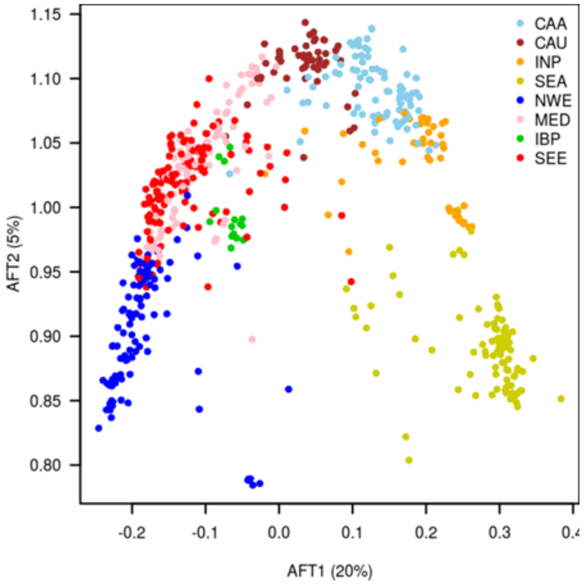


**Fig. S7. PCoA calculated with 8,741 haplotypes on 632 landraces.**

Simple Matching distance was calculated using Darwin software.
Cyan: Central Asia & Africa (CAA); Brown: Caucasus (CAU); Orange: Indian Peninsula (INP); Yellow: South-East Asia (SEA); Blue: Nort West Europe (NWE); Pink: Mediterranean basin (MED); Green: Iberian Peninsula (IBP); Red: South East Europe (SEE).

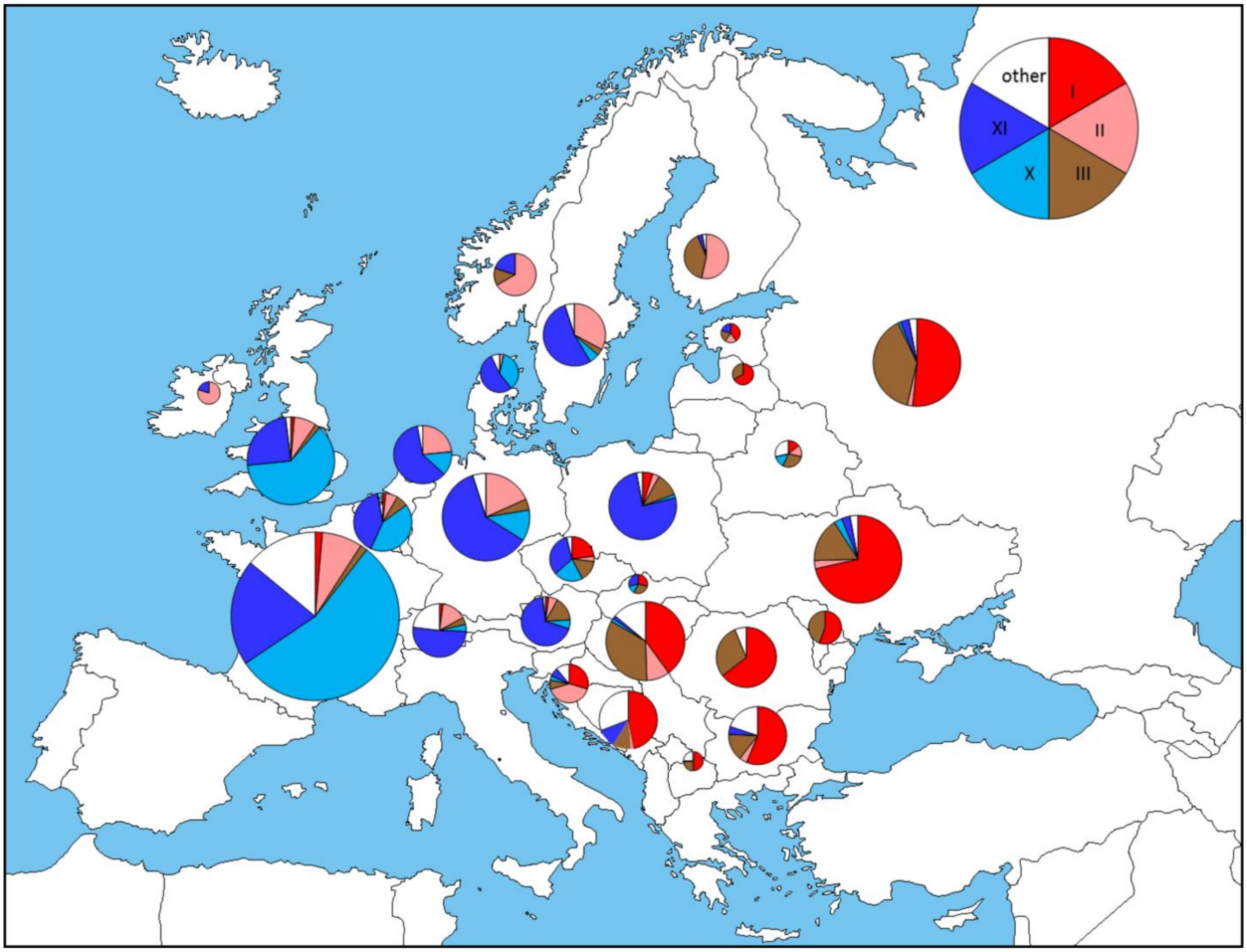


**Fig. S8. Frequency of major haplotype groups in Europe.**

The size of a pie is proportional to the number of accessions in each country.

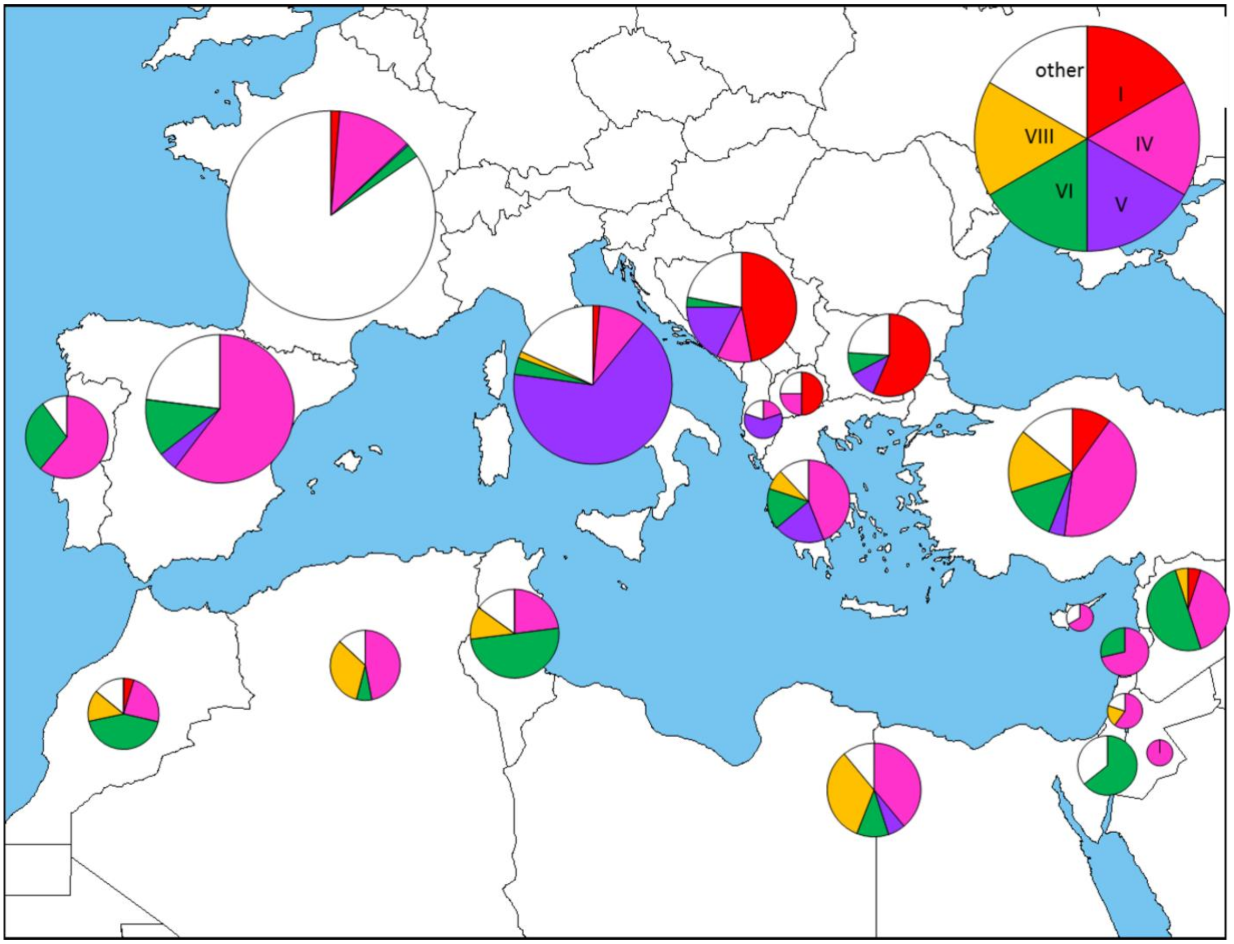


**Fig. S9. Frequency of major haplotype groups in the Mediterranean Basin.**
The size of a pie is proportional to the number of accessions in each country.

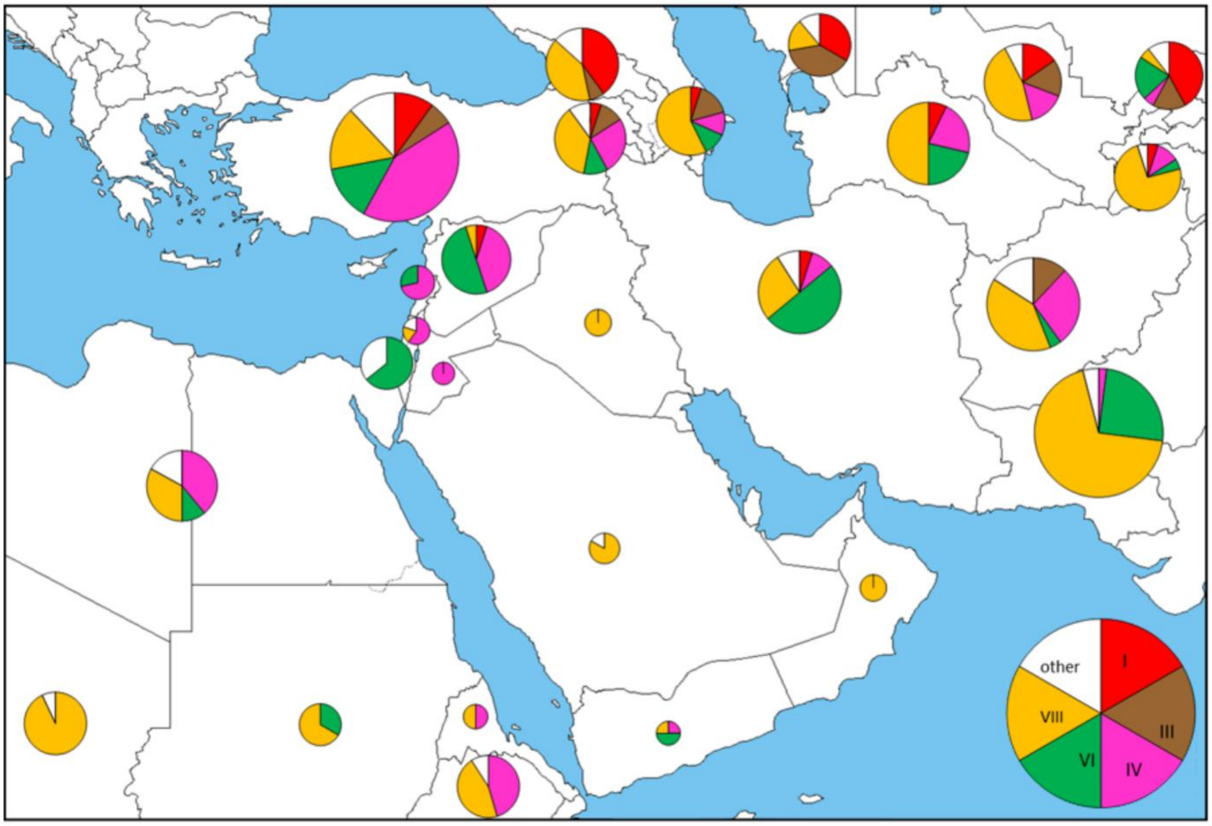


**Fig. S10. Frequency of major haplotype groups in Middle East and Central Asia.**
The size of a pie is proportional to the number of accessions in each country.

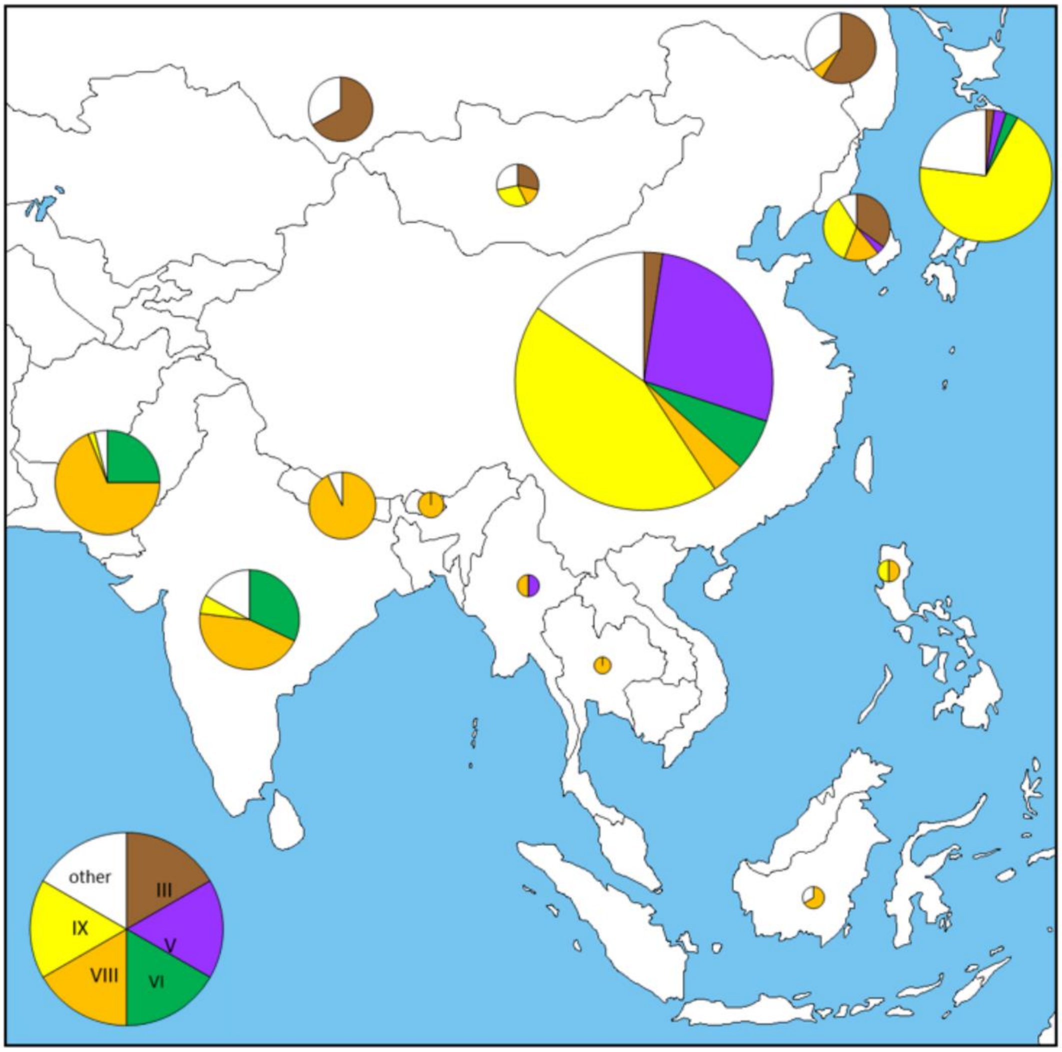


**Fig. S11. Frequency of major haplotype groups in South Eastern Asia.**
The size of a pie is proportional to the number of accessions in each country.

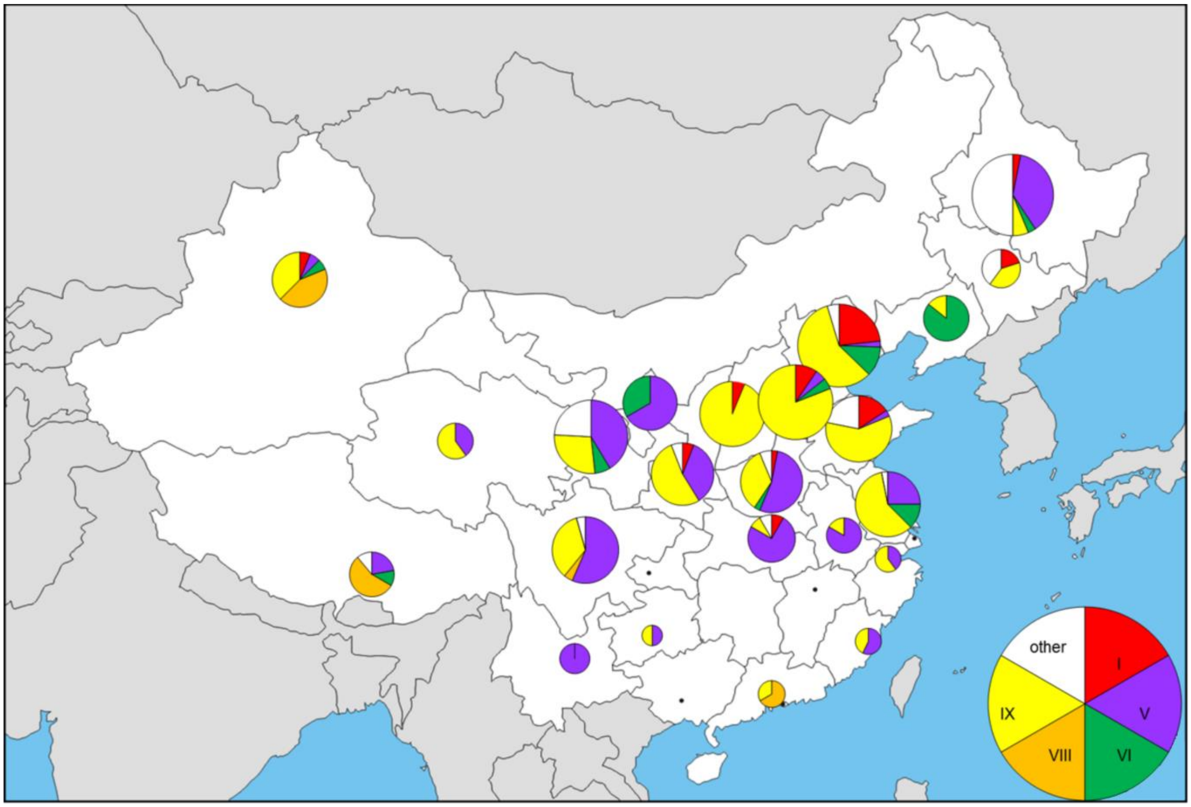


**Fig. S12. Frequency of major haplotype groups in China.**
The size of a pie is proportional to the number of accessions in each country.

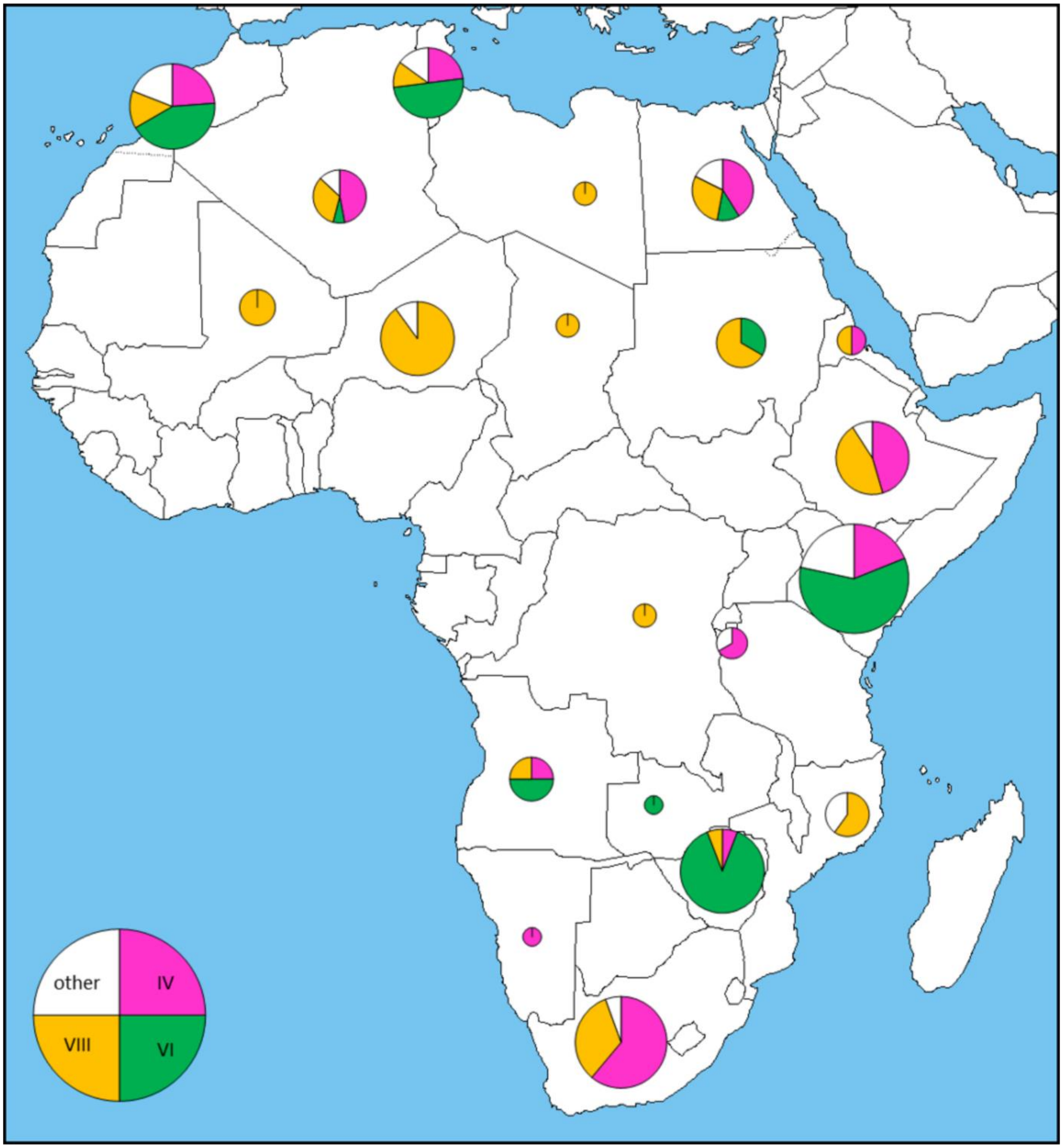


**Fig. S13. Frequency of major haplotype groups in Africa.**

The size of a pie is proportional to the number of accessions in each country.

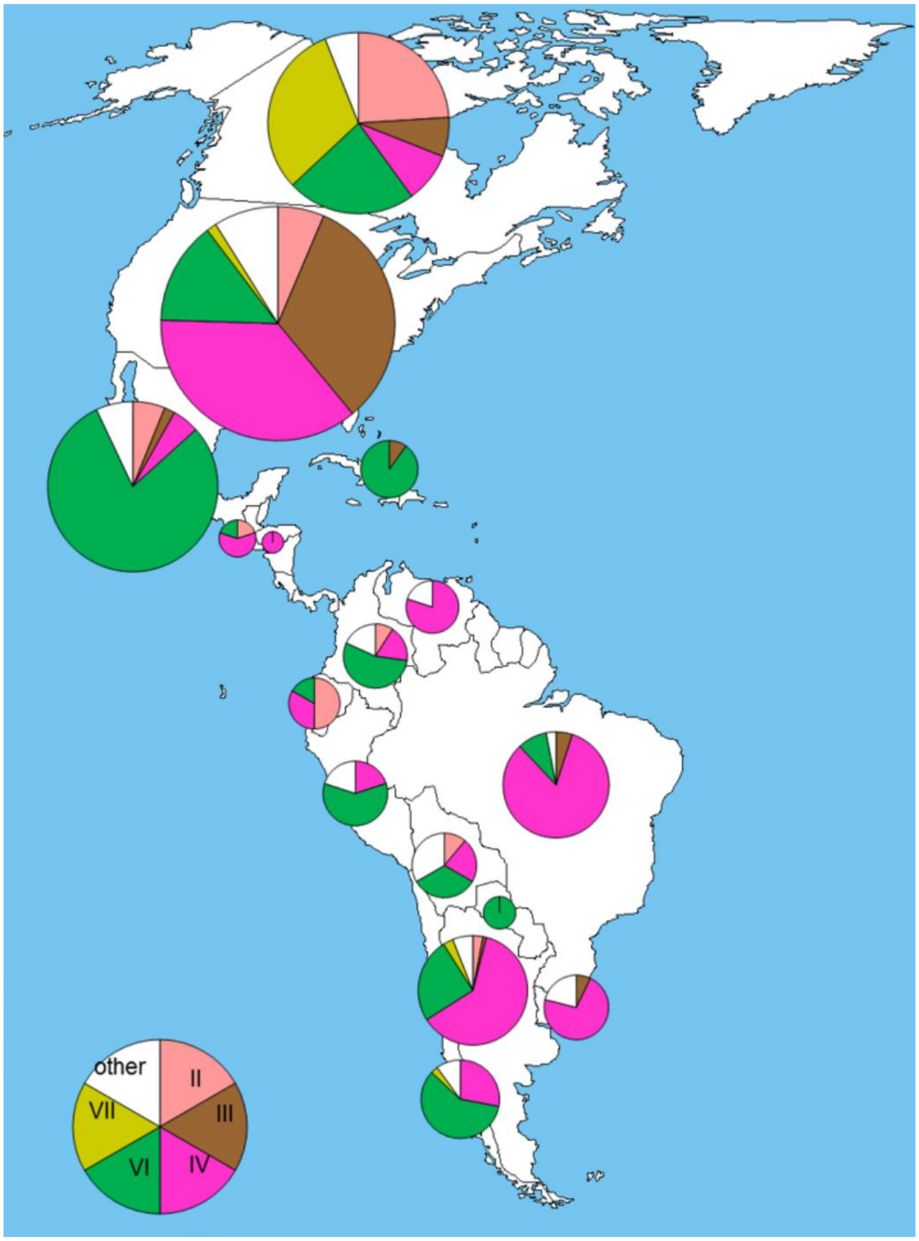


**Fig. S14. Frequency of major haplotype groups in America.**

The size of a pie is proportional to the number of accessions in each country.

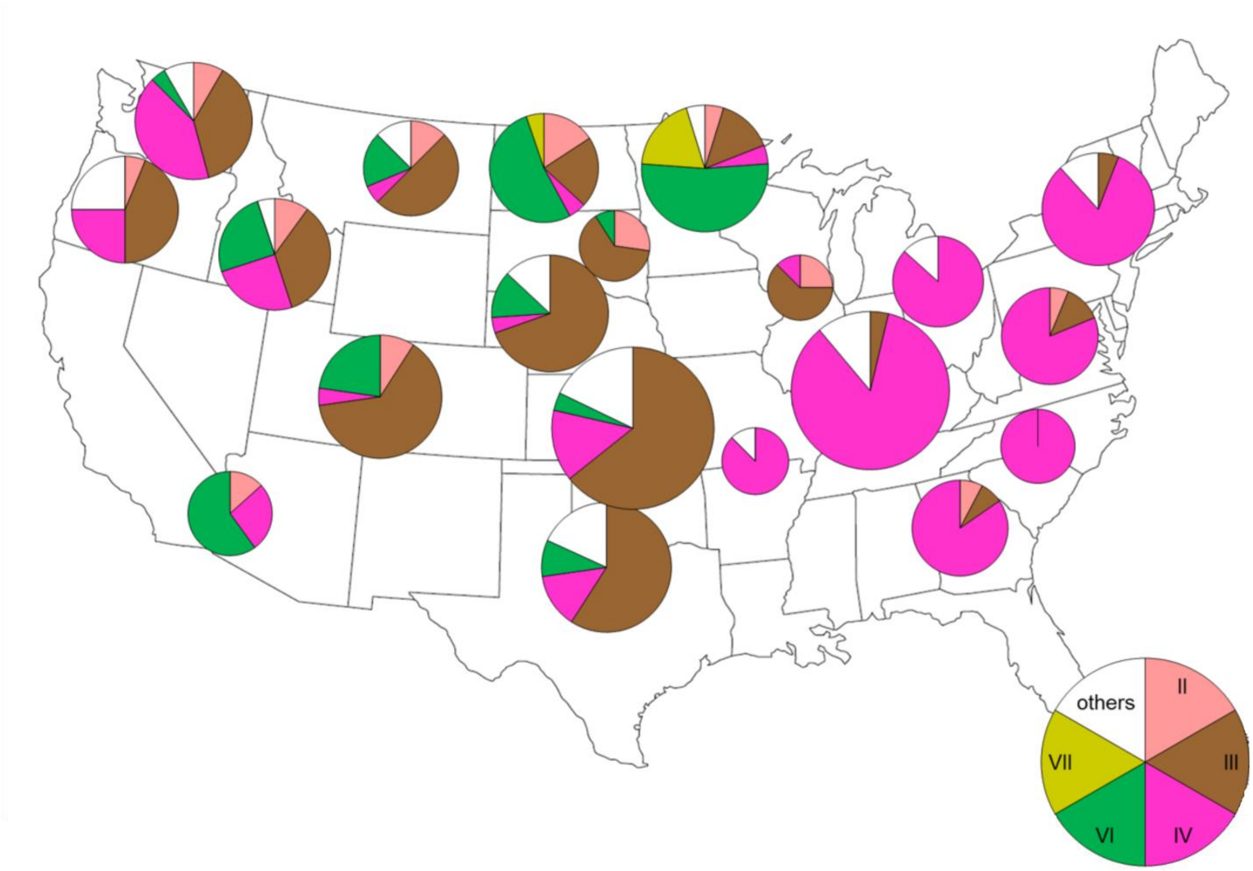


**Fig. S15. Frequency of major haplotype groups in the USA.**

The size of a pie is proportional to the number of accessions in each country.

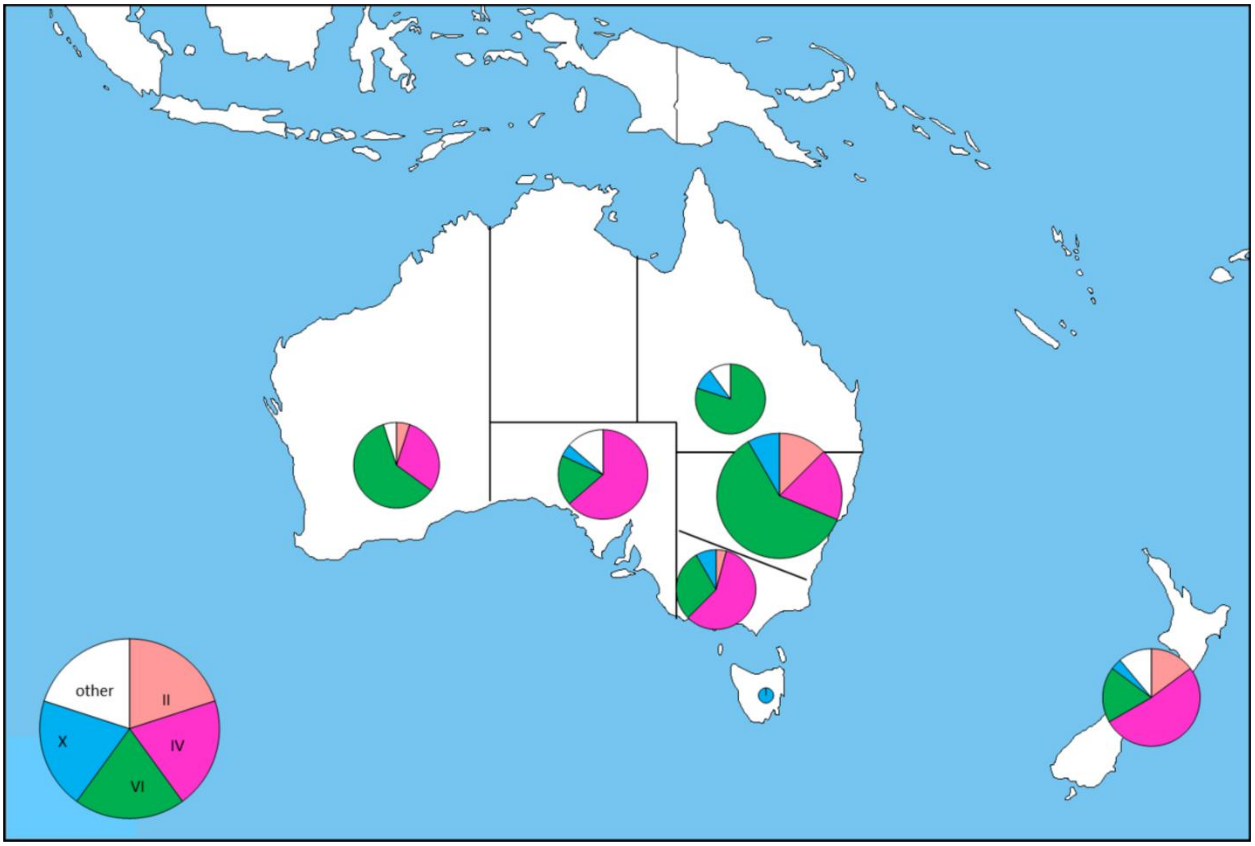


**Fig. S16. Frequency of major haplotype groups in Oceania.**

The size of a pie is proportional to the number of accessions in each country.

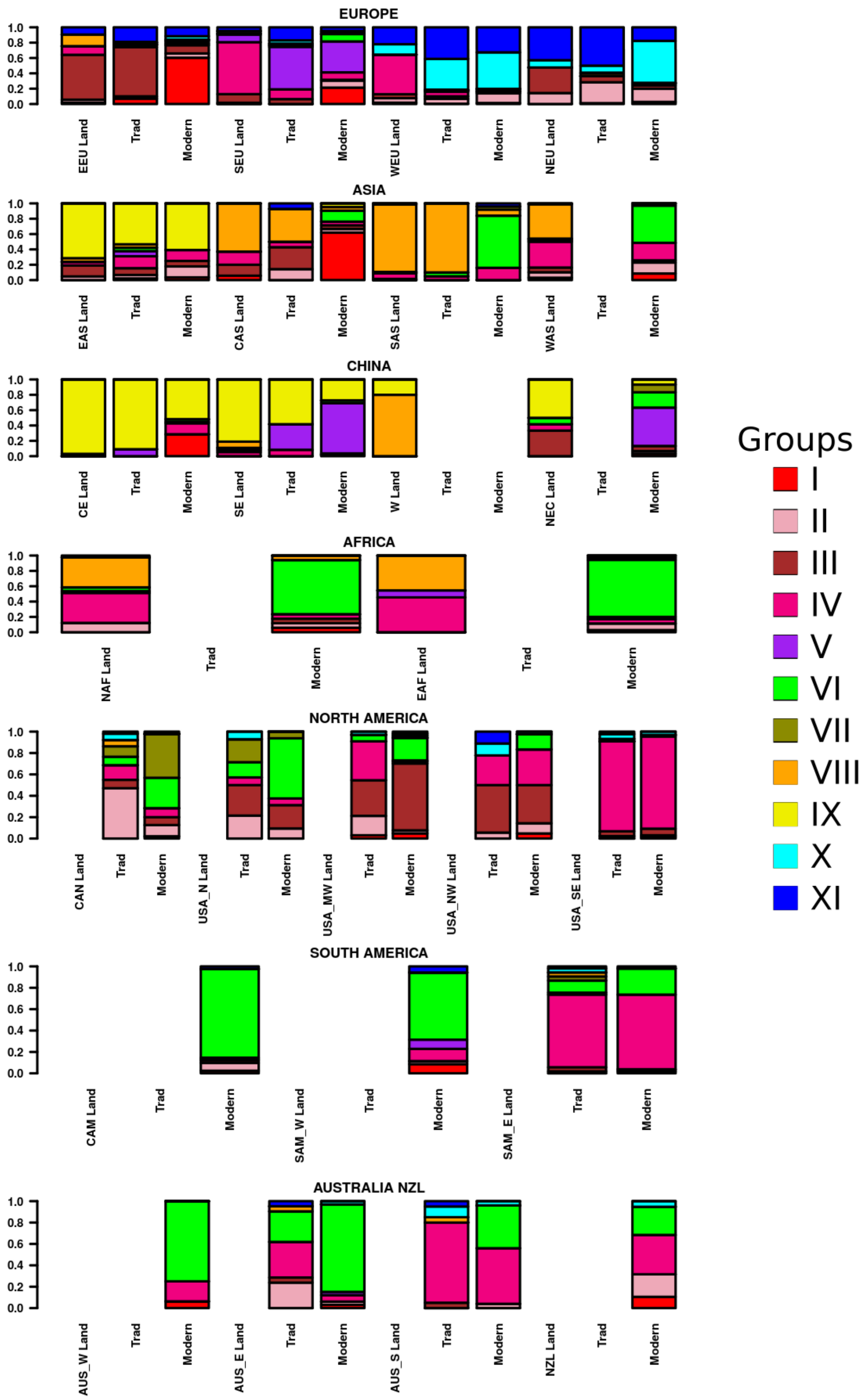
EUROPE
EEU Land
Trad
Modern
SEU Land
Trad
Modern
WEU Land
Trad
Modern
NEU Land
Trad
Modern
ASIA
EAS Land
Trad
Modern
CAS Land
Trad
Modern
SAS Land
Trad
Modern
WAS Land
Trad
Modern
CHINA
CE Land
Trad
Modern
SE Land
Trad
Modern
W Land
Trad
Modern
NEC Land
Trad
Modern
AFRICA
NAF Land
Trad
Modern
EAF Land
Trad
Modern
NORTH AMERICA
CAN Land
Trad
Modern
USA_N Land
Trad
Modern
USA_MW Land
Trad
Modern
USA_NW Land
Trad
Modern
USA_SE Land
Trad
Modern
SOUTH AMERICA
CAM Land
Trad
Modern
SAM_W Land
Trad
Modern
SAM_E Land
Trad
Modern
AUSTRALIA NZL
AUS_W Land
Trad
Modern
AUS_E Land
Trad
Modern
AUS_S Land
Trad
Modern
NZL Land
Trad
Modern
1.0
0.8
0.6
0.4
0.2
0.0
Groups
I
II
III
IV
V
VI
VII
VIII
IX
X
XI

**Fig. S17. Proportion of the eleven groups among the landraces, traditional cultivars and modern varieties from different region of the World.**

EUROPE - EEU: Eastern Europe; NEU: Northern Europe; SEU: Southern Europe; WEU: Western Europe
ASIA - CAS: Central Asia; EAS: Eastern Asia; SAS: Southern Asia; WAS: Western Asia
CHINA - SE: South-Eastern; W: Western; NEC: North-Eastern and Central
AFRICA - NAF: Northern Africa; EAF : Eastern Africa
NORTH AMERICA - CAN: Canada; USA_N: Northern USA; USA_MW: Mid-Western USA; USA_SE: Southern and Eastern USA, USA_NW: North-Western USA
SOUTH AMERICA - CAM: Central America, SAM_E: Eastern Coast of Southern America; SAM_W: Western Coast of Southern America
ASUTRALIA_NZL - AUS_W: Western Coast Australia; AUS_E: Eastern Coast Australia; AUS_S: Southern Coast Australia; NZL: New-Zealand

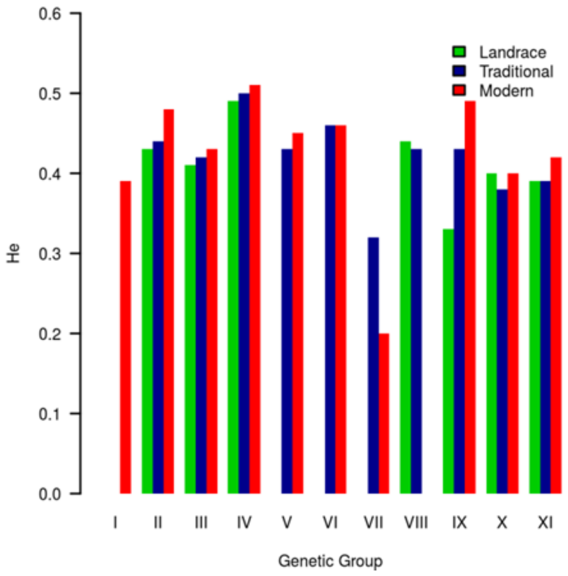


**Fig. S18. Nei Diversity level ( He) in the eleven haplotypic groups, according to the status of the accession (landraces, traditional and modern lines).**

**Table S1. Description of the eight sub-populations of landraces**

| Group[a] | N[b] | Assign[c] | Countries[d] | Regions[e] | Winter[f] | He[g] | Poly Rate[h] |
|---|---|---|---|---|---|---|---|
| CAA | 87 (54) | 68 (65) | OMN (9),SDN (7),SAU (7),PAK (7),IRN (7) | WAS (30),SAS (22),NAF (20),WAF (11),CAS (9) | 14 | 0.41 | 0.80 |
| CAU | 50 (38) | 80 (95) | AZE (24),RUS (19),TUR (16),GEO (14),ARM (11) | WAS (68),EEU (22),SEU (3),SEA (3),EAS (3) | 35 | 0.40 | 0.75 |
| INP | 47 (36) | 77 (73) | IND (28),PAK (25),NPL (22),IRQ (6),EGY (6) | SAS (83),WAS (6),SEA (6),NAF (6) | 0 | 0.35 | 0.71 |
| SEA | 93 (80) | 88 (99) | CHN (85),JPN (10) | EAS (98) | 53 | 0.34 | 0.77 |
| NWE | 107 (68) | 68 (66) | FRA (46),DEU (13),GBR (12),ESP (4),TUR (3) | WEU (60),NEU (15),SEU (12),EEU (6),WAS (4) | 84 | 0.45 | 0.88 |
| MED | 103 (59) | 66 (67) | SYR (12),FRA (10),ESP (10),ITA (7),TUR (5) | WAS (29),SEU (27),NAF (17),WEU (10),SAS (3) | 15 | 0.47 | 0.90 |
| IBP | 15 (12) | 89 (100) | ESP (75),PRT (25) | SEU (100) | 25 | 0.30 | 0.51 |
| SEE | 130 (84) | 71 (71) | FRA (14),RUS (12),HUN (11),UKR (5),CHN (5) | EEU (35),WEU (14),WAS (13),NEU (11),SEU (7) | 51 | 0.44 | 0.92 |

a Groups of landraces as defined by STRUCTURE algorithm. CAA: Central Asia – Africa; CAU: Caucasus; IBP: Iberian Peninsula; INP: Indian Peninsula; NWE: North Western Europe; MED: Mediterranean region; SEA: South Eastern Asia; SEE: South Eastern Europe

b Number of accessions with highest genome proportion (with at least 60% of its genome ) assigned to this group.

c Mean (median) genome proportion assigned to this group.

d Percentage of accessions originating from each country (International Organization for Standardization code). ARM: Armenia; AZE: Azerbaijan; CHN: China; DEU: Germany; EGY: Egypt; ESP: Spain; FRA: France; GBR: United Kingdom of Great Britain and Northern Ireland; GEO: Georgia; HUN: Hungary; IND: India; IRN: Islamic Republic of Iran; IRQ: Iraq; ITA: Italy; JPN: Japan; NPL: Nepal; OMN:Oman; PAK:Pakistan; PRT: Portugal; RUS: Russian Federation; SAU: Saudi Arabia; SDN: Sudan; SYR: Syrian Arab Republic; TUR: Turkey; UKR: Ukraine

e Percentage of accessions originating from each region (ISO code). CAS: Central Asia; EAS: Eastern Asia; EEU: Eastern Europe; NAF: Northern Africa; NEU: Northern Europe; SAS: Southern Asia; SEA: South-Eastern Asia; SEU: Southern Europe; WAF: Western Africa; WAS: Western Asia; WEU: Western Europe

f Percentage of winter type accessions. Most of other accessions are spring type, a few are alternative type.
g Nei He diversity index
h Percentage of polymorphic alleles

**Table S2. Pairwise differentiation index (Fst) matrix between the eight sub-populations of landraces**

| | CAA | CAU | INP | SEA | NWE | MED | IBP | SEE |
|---|---|---|---|---|---|---|---|---|
| CAA | | 0.13 | 0.17 | 0.21 | 0.25 | 0.17 | 0.36 | 0.20 |
| CAU | 0.13 | | 0.25 | 0.30 | 0.22 | 0.14 | 0.37 | 0.16 |
| INP | 0.17 | 0.25 | | 0.25 | 0.32 | 0.27 | 0.45 | 0.28 |
| SEA | 0.21 | 0.30 | 0.25 | | 0.37 | 0.32 | 0.47 | 0.33 |
| NWE | 0.25 | 0.22 | 0.32 | 0.37 | | 0.10 | 0.31 | 0.10 |
| MED | 0.17 | 0.14 | 0.27 | 0.32 | 0.10 | | 0.26 | 0.09 |
| IBP | 0.36 | 0.37 | 0.45 | 0.47 | 0.31 | 0.26 | | 0.31 |
| SEE | 0.20 | 0.16 | 0.28 | 0.33 | 0.10 | 0.09 | 0.31 | |

CAA: Central Asia – Africa; CAU: Caucasus; IBP: Iberian Peninsula; INP: Indian Peninsula; NWE: North Western Europe; MED: Mediterranean region; SEA: South Eastern Asia; SEE: South Eastern Europe. Lowest Fst values in red; highest Fst values in green.

#### Table S3. Description of the eleven groups of accessions

| Gp[a] | Name | N[b] | Countries[c] | Regions[d] | Landraces[e] | Registration[f] | Modern[g] | Winter[h] | He[i] | Poly[j] |
|---|---|---|---|---|---|---|---|---|---|---|
| I | SEE_modern | 413 | UKR (17),HUN (14),RUS (13),YUG (8),ROU (8) | EEU (64),SEU (11),EAS (7),CAS (4),WAS (3) | SEE (7) | 1984 (1984) | 95 | 93 | 0.39 | 0.95 |
| II | NWE_NAM | 353 | FRA (14),CAN (10),DEU (8),USA (7),SWE (5) | WEU (30),NEU (17),NAM (16),EEU (9),SEU (7) | SEE (19) | 1962 (1970) | 66 | 20 | 0.47 | 0.97 |
| III | NAM_SEE | 419 | USA (28),RUS (14),HUN (12),UKR (4),ROU (3) | EEU (40),NAM (30),WEU (7),NEU (6),EAS (5) | SEE (69) | 1960 (1965) | 57 | 86 | 0.43 | 0.97 |
| IV | NW_MED | 674 | USA (20),FRA (11),ARG (6),AUS (6),ESP (4) | NAM (22),SAM (15),WEU (14),SEU (12),AUS (8) | IBP (100),MED (100),NWE (9),CAU (8),SEE (5) | 1956 (1962) | 54 | 39 | 0.51 | 0.99 |
| V | CHN_IT_modern | 265 | CHN (40),ITA (37),YUG (5),HUN (4),GRC (2) | SEU (45),EAS (42),EEU (6) | NWE (10) | 1967 (1968) | 81 | 39 | 0.45 | 0.94 |
| VI | CIMMYT_NW_CHN_modern | 528 | MEX (15),AUS (11),USA (10),CAN (6),CHN (5) | NAM (16),CAM (15),AUS (12),SAM (11),SAS (8) | - | 1976 (1977) | 90 | 7 | 0.47 | 0.97 |
| VII | CAN_CHN_USA_modern | 63 | CAN (70),CHN (11),USA (8),ARG (3) | NAM (78),EAS (11),SAM (5) | - | 1971 (1974) | 77 | 0 | 0.24 | 0.72 |
| VIII | SEA_INP | 292 | PAK (11),NPL (9),IND (8),CHN (5),TJK (5) | SAS (35),WAS (17),CAS (11),NAF (7),EAS (7) | INP (100),CAA (100),CAU (92) | 1938 (1934) | 16 | 23 | 0.44 | 0.91 |
| IX | CHN_JPN | 255 | CHN (66),JPN (23),KOR (3) | EAS (93) | SEA (98) | 1961 (1965) | 63 | 63 | 0.45 | 0.93 |
| X | FRA_GBR_modern | 581 | FRA (62),GBR (17),BEL (4),DEU (3) | WEU (72),NEU (19),EEU (4) | NWE (24) | 1969 (1974) | 73 | 90 | 0.40 | 0.94 |
| XI | FRA_DEU | 560 | FRA (24),DEU (16),POL (8),GBR (7),NLD (6) | WEU (61),NEU (16),EEU (14),SEU (4) | NWE (56) | 1957 (1964) | 57 | 96 | 0.42 | 0.94 |

a Haplotypic Groups, from I to XI

b Number of accessions

c Percentage of accessions originating from each country (ISO code). ARG: Argentina; AUS: Australia; BEL: Belgium; CAN: Canada; CHN: China; DEU: Germany; ESP: Spain; FRA: France; GBR: United Kingdom of Great Britain and Northern Ireland; GRC: Greece; HUN: Hungary; IND: India; ITA: Italy; JPN: Japan; KOR: Republic of Korea; MEX: Mexico; NLD: Netherlands; NPL: Nepal; PAK: Pakistan; POL: Poland; ROU: Romania; RUS: Russian Federation; SWE: Sweden; TJK: Tajikistan; UKR: Ukraine; USA: United States of America; YUG: Yugoslavia d Percentage of accessions originating from each region (ISO code). AUS: Australia and NewZealand; CAM: Central America; CAS: Central Asia; EAS: Eastern Asia; EEU: Eastern Europe; NAF: Northern Africa; NAM: Northern America; NEU: Northern Europe; SAM: South America; SAS: Southern Asia; SEU: Southern Europe; WAS: Western Asia; WEU: Western Europe

e Percentage of accessions from the eight landrace groups defined by STRUCTURE assigned to this group

f Mean (Median) registration date

g Percentage of accessions registered after 1959

h Percentage of winter type accessions. Most of other accessions are spring type, a few are alternative type

i Nei He diversity index

j Percentage of polymorphic alleles

**Table S4. Pairwise differentiation index (Fst) matrix between the eleven groups (4,403 accessions)**

| | I | II | III | IV | V | VI | VII | VIII | IX | X | XI |
|---|---|---|---|---|---|---|---|---|---|---|---|
| I | | 0.15 | 0.11 | 0.12 | 0.19 | 0.17 | 0.37 | 0.27 | 0.27 | 0.23 | 0.19 |
| II | 0.15 | | 0.07 | 0.05 | 0.13 | 0.09 | 0.25 | 0.17 | 0.19 | 0.15 | 0.12 |
| III | 0.11 | 0.07 | | 0.06 | 0.17 | 0.13 | 0.28 | 0.19 | 0.21 | 0.20 | 0.15 |
| IV | 0.12 | 0.05 | 0.06 | | 0.10 | 0.07 | 0.25 | 0.14 | 0.16 | 0.13 | 0.11 |
| V | 0.19 | 0.13 | 0.17 | 0.10 | | 0.13 | 0.35 | 0.22 | 0.19 | 0.16 | 0.14 |
| VI | 0.17 | 0.09 | 0.13 | 0.07 | 0.13 | | 0.24 | 0.18 | 0.19 | 0.21 | 0.18 |
| VII | 0.37 | 0.25 | 0.28 | 0.25 | 0.35 | 0.24 | | 0.35 | 0.37 | 0.39 | 0.37 |
| VIII | 0.27 | 0.17 | 0.19 | 0.14 | 0.22 | 0.18 | 0.35 | | 0.11 | 0.29 | 0.27 |
| IX | 0.27 | 0.19 | 0.21 | 0.16 | 0.19 | 0.19 | 0.37 | 0.11 | | 0.31 | 0.28 |
| X | 0.23 | 0.15 | 0.20 | 0.13 | 0.16 | 0.21 | 0.39 | 0.29 | 0.31 | | 0.07 |
| XI | 0.19 | 0.12 | 0.15 | 0.11 | 0.14 | 0.18 | 0.37 | 0.27 | 0.28 | 0.07 | |

### Table S5. Pairwise differentiation index (Fst) matrix between the eight sub-populations of landraces and the eleven groups

| | VII_trad | VII_mod | IBP | SEA | CAA | INP | VIII_trad | VIII_mod | IX_trad | IX_mod | CAU | I_trad | I_mod | VI_trad | VI_mod | MED | SEE | III_trad | III_mod | II_trad | II_mod | IV_trad | IV_mod | NWE | XI_trad | XI_mod | X_trad | X_mod | V_trad | V_mod |
|---|---|---|---|---|---|---|---|---|---|---|---|---|---|---|---|---|---|---|---|---|---|---|---|---|---|---|---|---|---|---|
| VII_trad | NA | 0.09 | 0.44 | 0.44 | 0.32 | 0.40 | 0.29 | 0.29 | 0.32 | 0.24 | 0.31 | 0.35 | 0.31 | 0.18 | 0.18 | 0.23 | 0.21 | 0.23 | 0.20 | 0.17 | 0.18 | 0.19 | 0.17 | 0.27 | 0.34 | 0.30 | 0.35 | 0.34 | 0.31 | 0.27 |
| VII_mod | 0.09 | NA | 0.59 | 0.54 | 0.44 | 0.52 | 0.40 | 0.45 | 0.46 | 0.35 | 0.45 | 0.52 | 0.39 | 0.31 | 0.26 | 0.35 | 0.33 | 0.34 | 0.31 | 0.30 | 0.28 | 0.30 | 0.27 | 0.39 | 0.43 | 0.39 | 0.45 | 0.42 | 0.45 | 0.37 |
| IBP | 0.44 | 0.59 | NA | 0.47 | 0.36 | 0.45 | 0.34 | 0.37 | 0.37 | 0.30 | 0.37 | 0.46 | 0.38 | 0.29 | 0.29 | 0.26 | 0.31 | 0.33 | 0.32 | 0.33 | 0.28 | 0.21 | 0.22 | 0.31 | 0.38 | 0.35 | 0.39 | 0.38 | 0.36 | 0.32 |
| SEA | 0.44 | 0.54 | 0.47 | NA | 0.21 | 0.25 | 0.18 | 0.21 | 0.06 | 0.13 | 0.30 | 0.45 | 0.40 | 0.33 | 0.30 | 0.32 | 0.33 | 0.35 | 0.33 | 0.35 | 0.31 | 0.29 | 0.27 | 0.37 | 0.42 | 0.40 | 0.44 | 0.42 | 0.33 | 0.31 |
| CAA | 0.32 | 0.44 | 0.36 | 0.21 | NA | 0.17 | 0.05 | 0.08 | 0.15 | 0.13 | 0.13 | 0.31 | 0.30 | 0.21 | 0.21 | 0.17 | 0.20 | 0.23 | 0.21 | 0.22 | 0.20 | 0.17 | 0.16 | 0.25 | 0.32 | 0.29 | 0.33 | 0.32 | 0.26 | 0.23 |
| INP | 0.40 | 0.52 | 0.45 | 0.25 | 0.17 | NA | 0.05 | 0.04 | 0.20 | 0.19 | 0.25 | 0.41 | 0.36 | 0.27 | 0.26 | 0.27 | 0.28 | 0.30 | 0.29 | 0.28 | 0.26 | 0.24 | 0.23 | 0.32 | 0.38 | 0.35 | 0.40 | 0.38 | 0.32 | 0.29 |
| VIII_trad | 0.29 | 0.40 | 0.34 | 0.18 | 0.05 | 0.05 | NA | 0.01 | 0.12 | 0.11 | 0.11 | 0.28 | 0.28 | 0.18 | 0.18 | 0.16 | 0.18 | 0.21 | 0.19 | 0.19 | 0.17 | 0.15 | 0.14 | 0.23 | 0.30 | 0.27 | 0.31 | 0.31 | 0.23 | 0.21 |
| VIII_mod | 0.29 | 0.45 | 0.37 | 0.21 | 0.08 | 0.04 | 0.01 | NA | 0.14 | 0.11 | 0.14 | 0.30 | 0.29 | 0.18 | 0.18 | 0.17 | 0.18 | 0.21 | 0.19 | 0.18 | 0.17 | 0.16 | 0.15 | 0.24 | 0.31 | 0.28 | 0.33 | 0.32 | 0.24 | 0.22 |
| IX_trad | 0.32 | 0.46 | 0.37 | 0.06 | 0.15 | 0.20 | 0.12 | 0.14 | NA | 0.03 | 0.20 | 0.31 | 0.30 | 0.20 | 0.20 | 0.21 | 0.22 | 0.24 | 0.22 | 0.23 | 0.20 | 0.18 | 0.17 | 0.25 | 0.32 | 0.29 | 0.34 | 0.33 | 0.20 | 0.20 |
| IX_mod | 0.24 | 0.35 | 0.30 | 0.13 | 0.13 | 0.19 | 0.11 | 0.11 | 0.03 | NA | 0.15 | 0.20 | 0.20 | 0.13 | 0.13 | 0.13 | 0.13 | 0.15 | 0.12 | 0.15 | 0.12 | 0.11 | 0.09 | 0.16 | 0.23 | 0.21 | 0.25 | 0.25 | 0.13 | 0.13 |
| CAU | 0.31 | 0.45 | 0.37 | 0.30 | 0.13 | 0.25 | 0.11 | 0.14 | 0.20 | 0.15 | NA | 0.29 | 0.27 | 0.20 | 0.19 | 0.14 | 0.16 | 0.18 | 0.17 | 0.19 | 0.17 | 0.14 | 0.14 | 0.22 | 0.29 | 0.27 | 0.31 | 0.30 | 0.25 | 0.22 |
| I_trad | 0.35 | 0.52 | 0.46 | 0.45 | 0.31 | 0.41 | 0.28 | 0.30 | 0.31 | 0.20 | 0.29 | NA | 0.07 | 0.21 | 0.20 | 0.19 | 0.11 | 0.11 | 0.11 | 0.19 | 0.15 | 0.15 | 0.13 | 0.18 | 0.24 | 0.22 | 0.28 | 0.27 | 0.26 | 0.22 |
| I_mod | 0.31 | 0.39 | 0.38 | 0.40 | 0.30 | 0.36 | 0.28 | 0.29 | 0.30 | 0.20 | 0.27 | 0.07 | NA | 0.20 | 0.17 | 0.18 | 0.13 | 0.13 | 0.11 | 0.19 | 0.14 | 0.14 | 0.12 | 0.16 | 0.21 | 0.19 | 0.24 | 0.23 | 0.22 | 0.19 |
| VI_trad | 0.18 | 0.31 | 0.29 | 0.33 | 0.21 | 0.27 | 0.18 | 0.18 | 0.20 | 0.13 | 0.20 | 0.21 | 0.20 | NA | 0.02 | 0.10 | 0.11 | 0.14 | 0.12 | 0.11 | 0.08 | 0.08 | 0.06 | 0.14 | 0.21 | 0.19 | 0.23 | 0.23 | 0.14 | 0.13 |
| VI_mod | 0.18 | 0.26 | 0.29 | 0.30 | 0.21 | 0.26 | 0.18 | 0.18 | 0.20 | 0.13 | 0.19 | 0.20 | 0.17 | 0.02 | NA | 0.11 | 0.12 | 0.14 | 0.12 | 0.12 | 0.09 | 0.09 | 0.06 | 0.14 | 0.20 | 0.18 | 0.21 | 0.21 | 0.14 | 0.13 |
| MED | 0.23 | 0.35 | 0.26 | 0.32 | 0.17 | 0.27 | 0.16 | 0.17 | 0.21 | 0.13 | 0.14 | 0.19 | 0.18 | 0.10 | 0.11 | NA | 0.09 | 0.11 | 0.11 | 0.13 | 0.09 | 0.02 | 0.04 | 0.10 | 0.17 | 0.16 | 0.17 | 0.19 | 0.15 | 0.13 |
| SEE | 0.21 | 0.33 | 0.31 | 0.33 | 0.20 | 0.28 | 0.18 | 0.18 | 0.22 | 0.13 | 0.16 | 0.11 | 0.13 | 0.11 | 0.12 | 0.09 | NA | 0.01 | 0.03 | 0.05 | 0.05 | 0.06 | 0.05 | 0.10 | 0.16 | 0.15 | 0.20 | 0.20 | 0.18 | 0.15 |
| III_trad | 0.23 | 0.34 | 0.33 | 0.35 | 0.23 | 0.30 | 0.21 | 0.21 | 0.24 | 0.15 | 0.18 | 0.11 | 0.13 | 0.14 | 0.14 | 0.11 | 0.01 | NA | 0.02 | 0.08 | 0.08 | 0.08 | 0.07 | 0.12 | 0.18 | 0.16 | 0.22 | 0.21 | 0.20 | 0.17 |
| III_mod | 0.20 | 0.31 | 0.32 | 0.33 | 0.21 | 0.29 | 0.19 | 0.19 | 0.22 | 0.12 | 0.17 | 0.11 | 0.11 | 0.12 | 0.12 | 0.11 | 0.03 | 0.02 | NA | 0.09 | 0.07 | 0.07 | 0.06 | 0.12 | 0.18 | 0.16 | 0.21 | 0.21 | 0.18 | 0.15 |
| II_trad | 0.17 | 0.30 | 0.33 | 0.35 | 0.22 | 0.28 | 0.19 | 0.18 | 0.23 | 0.15 | 0.19 | 0.19 | 0.19 | 0.11 | 0.12 | 0.13 | 0.05 | 0.08 | 0.09 | NA | 0.04 | 0.09 | 0.08 | 0.15 | 0.20 | 0.18 | 0.23 | 0.23 | 0.20 | 0.17 |
| II_mod | 0.18 | 0.28 | 0.28 | 0.31 | 0.20 | 0.26 | 0.17 | 0.17 | 0.20 | 0.12 | 0.17 | 0.15 | 0.14 | 0.08 | 0.09 | 0.09 | 0.05 | 0.08 | 0.07 | 0.04 | NA | 0.05 | 0.04 | 0.07 | 0.12 | 0.10 | 0.13 | 0.14 | 0.13 | 0.10 |
| IV_trad | 0.19 | 0.30 | 0.21 | 0.29 | 0.17 | 0.24 | 0.15 | 0.16 | 0.18 | 0.11 | 0.14 | 0.15 | 0.14 | 0.08 | 0.09 | 0.02 | 0.06 | 0.08 | 0.07 | 0.09 | 0.05 | NA | 0.01 | 0.06 | 0.12 | 0.12 | 0.14 | 0.15 | 0.12 | 0.10 |
| IV_mod | 0.17 | 0.27 | 0.22 | 0.27 | 0.16 | 0.23 | 0.14 | 0.15 | 0.17 | 0.09 | 0.14 | 0.13 | 0.12 | 0.06 | 0.06 | 0.04 | 0.05 | 0.07 | 0.06 | 0.08 | 0.04 | 0.01 | NA | 0.07 | 0.13 | 0.11 | 0.14 | 0.15 | 0.11 | 0.09 |
| NWE | 0.27 | 0.39 | 0.31 | 0.37 | 0.25 | 0.32 | 0.23 | 0.24 | 0.25 | 0.16 | 0.22 | 0.18 | 0.16 | 0.14 | 0.14 | 0.10 | 0.10 | 0.12 | 0.12 | 0.15 | 0.07 | 0.06 | 0.07 | NA | 0.03 | 0.05 | 0.08 | 0.09 | 0.11 | 0.09 |
| XI_trad | 0.34 | 0.43 | 0.38 | 0.42 | 0.32 | 0.38 | 0.30 | 0.31 | 0.32 | 0.23 | 0.29 | 0.24 | 0.21 | 0.21 | 0.20 | 0.17 | 0.16 | 0.18 | 0.18 | 0.20 | 0.12 | 0.12 | 0.13 | 0.03 | NA | 0.03 | 0.09 | 0.10 | 0.18 | 0.15 |
| XI_mod | 0.30 | 0.39 | 0.35 | 0.40 | 0.29 | 0.35 | 0.27 | 0.28 | 0.29 | 0.21 | 0.27 | 0.22 | 0.19 | 0.19 | 0.18 | 0.16 | 0.15 | 0.16 | 0.16 | 0.18 | 0.10 | 0.12 | 0.11 | 0.05 | 0.03 | NA | 0.08 | 0.06 | 0.17 | 0.14 |
| X_trad | 0.35 | 0.45 | 0.39 | 0.44 | 0.33 | 0.40 | 0.31 | 0.33 | 0.34 | 0.25 | 0.31 | 0.28 | 0.24 | 0.23 | 0.21 | 0.17 | 0.20 | 0.22 | 0.21 | 0.23 | 0.13 | 0.14 | 0.14 | 0.08 | 0.09 | 0.08 | NA | 0.04 | 0.19 | 0.15 |
| X_mod | 0.34 | 0.42 | 0.38 | 0.42 | 0.32 | 0.38 | 0.31 | 0.32 | 0.33 | 0.25 | 0.30 | 0.27 | 0.23 | 0.23 | 0.21 | 0.19 | 0.20 | 0.21 | 0.21 | 0.23 | 0.14 | 0.15 | 0.15 | 0.09 | 0.10 | 0.06 | 0.04 | NA | 0.19 | 0.16 |
| V_trad | 0.31 | 0.45 | 0.36 | 0.33 | 0.26 | 0.32 | 0.23 | 0.24 | 0.20 | 0.13 | 0.25 | 0.26 | 0.22 | 0.14 | 0.14 | 0.15 | 0.18 | 0.20 | 0.18 | 0.20 | 0.13 | 0.12 | 0.11 | 0.11 | 0.18 | 0.17 | 0.19 | 0.19 | NA | 0.01 |
| V_mod | 0.27 | 0.37 | 0.32 | 0.31 | 0.23 | 0.29 | 0.21 | 0.22 | 0.20 | 0.13 | 0.22 | 0.22 | 0.19 | 0.13 | 0.13 | 0.13 | 0.15 | 0.17 | 0.15 | 0.17 | 0.10 | 0.10 | 0.09 | 0.09 | 0.15 | 0.14 | 0.15 | 0.16 | 0.01 | NA |

#### Table S6. Main structural variations (> 5Mb) detected in the wheat genome

| Chromosome | Start (Mb) | Stop (Mb) | Size (Mb) | Chromosome length (Mb) | Fraction of the chromosome |
|---|---|---|---|---|---|
| 1A | 0 | 214 | 214 | 593 | 36% |
| 1A | 0 | 28 | 28 | 593 | 5% |
| 1A | 1 | 22 | 21 | 593 | 4% |
| 1A | 565 | 593 | 28 | 593 | 5% |
| 1B | 0 | 689 | 689 | 689 | 100% |
| 1B | 0 | 236 | 236 | 689 | 34% |
| 1B | 0 | 218 | 218 | 689 | 32% |
| 1B | 625 | 689 | 64 | 689 | 9% |
| 1D | 0 | 12 | 12 | 495 | 2% |
| 1D | 413 | 418 | 5 | 495 | 1% |
| 1D | 413 | 424 | 11 | 495 | 2% |
| 2A | 0 | 10 | 10 | 780 | 1% |
| 2A | 13 | 19 | 6 | 780 | 1% |
| 2A | 611 | 780 | 169 | 780 | 22% |
| 2A | 712 | 780 | 68 | 780 | 9% |
| 2A | 734 | 780 | 46 | 780 | 6% |
| 2B | 90 | 757 | 667 | 800 | 83% |
| 2B | 54 | 733 | 679 | 800 | 85% |
| 2B | 90 | 609 | 519 | 800 | 65% |
| 2B | 90 | 154 | 64 | 800 | 8% |
| 2B | 455 | 733 | 278 | 800 | 35% |
| 2B | 591 | 768 | 177 | 800 | 22% |
| 2B | 670 | 743 | 73 | 800 | 9% |
| 2B | 0 | 30 | 30 | 800 | 4% |
| 2D | 0 | 18 | 18 | 651 | 3% |
| 2D | 0 | 46 | 46 | 651 | 7% |
| 2D | 0 | 457 | 457 | 651 | 70% |
| 2D | 636 | 651 | 15 | 651 | 2% |
| 3A | 0 | 41 | 41 | 751 | 5% |
| 3A | 0 | 19 | 19 | 751 | 3% |
| 3A | 8 | 14 | 6 | 751 | 1% |
| 3A | 732 | 751 | 19 | 751 | 3% |
| 3B | 7 | 14 | 7 | 829 | 1% |
| 3B | 1 | 33 | 32 | 829 | 4% |
| 3B | 1 | 6 | 5 | 829 | 1% |
| 3B | 0 | 123 | 123 | 829 | 15% |
| 3B | 819 | 829 | 10 | 829 | 1% |

#### Table S6. Main structural variations (> 5Mb) detected in the wheat genome (Cont.)

| Chromosome | Start (Mb) | Stop (Mb) | Size (Mb) | Chromosome length (Mb) | Fraction of the chromosome |
|---|---|---|---|---|---|
| 3D | 504 | 615 | 111 | 615 | 18% |
| 3D | 600 | 615 | 15 | 615 | 2% |
| 4A | 640 | 744 | 104 | 744 | 14% |
| 4A | 725 | 744 | 19 | 744 | 3% |
| 4A | 642 | 655 | 13 | 744 | 2% |
| 4B | 640 | 672 | 32 | 672 | 5% |
| 4B | 656 | 672 | 16 | 672 | 2% |
| 4D | 0 | 8 | 8 | 509 | 2% |
| 5A | 0 | 31 | 31 | 710 | 4% |
| 5A | 0 | 19 | 19 | 710 | 3% |
| 5A | 464 | 475 | 11 | 710 | 2% |
| 5A | 534 | 539 | 5 | 710 | 1% |
| 5B | 533 | 545 | 12 | 713 | 2% |
| 5B | 486 | 513 | 27 | 713 | 4% |
| 5B | 507 | 525 | 18 | 713 | 3% |
| 5B | 588 | 713 | 125 | 713 | 18% |
| 5D | 501 | 564 | 63 | 565 | 11% |
| 6A | 306 | 617 | 311 | 617 | 50% |
| 6A | 0 | 50 | 50 | 617 | 8% |
| 6A | 1 | 20 | 19 | 617 | 3% |
| 6A | 585 | 617 | 32 | 617 | 5% |
| 6B | 0 | 39 | 39 | 721 | 5% |
| 6B | 0 | 9 | 9 | 721 | 1% |
| 6B | 704 | 721 | 17 | 721 | 2% |
| 6D | 0 | 7 | 7 | 473 | 1% |
| 6D | 0 | 24 | 24 | 473 | 5% |
| 6D | 456 | 473 | 17 | 473 | 4% |
| 6D | 462 | 470 | 8 | 473 | 2% |
| 7A | 0 | 184 | 184 | 736 | 25% |
| 7A | 0 | 30 | 30 | 736 | 4% |
| 7A | 150 | 162 | 12 | 736 | 2% |
| 7A | 592 | 736 | 144 | 736 | 20% |
| 7A | 672 | 736 | 64 | 736 | 9% |
| 7A | 699 | 736 | 37 | 736 | 5% |

#### Table S6. Main structural variations (> 5Mb) detected in the wheat genome (Cont.)

| Chromosome | Start (Mb) | Stop (Mb) | Size (Mb) | Chromosome length (Mb) | Fraction of the chromosome |
|---|---|---|---|---|---|
| 7B | 0 | 73 | 73 | 751 | 10% |
| 7B | 34 | 40 | 6 | 751 | 1% |
| 7B | 633 | 751 | 118 | 751 | 16% |
| 7B | 668 | 751 | 83 | 751 | 11% |
| 7B | 727 | 732 | 5 | 751 | 1% |
| 7B | 693 | 698 | 5 | 751 | 1% |
| 7B | 739 | 749 | 10 | 751 | 1% |
| 7D | 0 | 10 | 10 | 639 | 2% |
| 7D | 49 | 59 | 10 | 639 | 2% |
| 7D | 0 | 457 | 457 | 639 | 72% |
| 7D | 375 | 639 | 264 | 639 | 41% |
| 7D | 563 | 639 | 76 | 639 | 12% |
| 7D | 612 | 639 | 27 | 639 | 4% |

**Data S1. (Separate file)**

List of 4,506 wheat accessions and related information.

**Data S2. (https://urgi.versailles.inra.fr/download/wheat/genotyping/Balfourier_et_al_Wheat_Phylogeography_DataS2.zip)**

Genotyping data of 4,506 wheat accession with 113,457 genome-wide SNPs.

**Data S3. (https://urgi.versailles.inra.fr/download/wheat/genotyping/Balfourier_et_al_Wheat_Phylogeography_DataS3.zip)**

Haplotyping data of 4,403 wheat accessions with 8,741 haplotypic blocks.

#### References

29. J. C. Barrett, B. Fry, J. Maller, M. J. Daly, Haploview: analysis and visualization of LD and haplotype maps. *Bioinformatics* **21**, 263 (2005).

30. S. Purcell *et al.*, PLINK: A Tool Set for Whole-Genome Association and Population-Based Linkage Analyses. *Am. J. Hum. Genet.* **81**, 559 (2007).

31. J. K. Pritchard, M. Stephens, P. Donnelly, Inference of population structure using multilocus genotype data. *Genetics* **155**, 945 (2000).

32. D. Falush, M. Stephens, J. K. Pritchard, Inference of population structure using multilocus genotype data: linked loci and correlated allele frequencies. *Genetics* **164**, 1567 (2003).

33. B. S. Weir, C. C. Cockerham, Estimating F-Statistics for the Analysis of Population Structure. *Evolution* **38**, 1358 (1984).

34. J. Goudet, hierfstat, a package for r to compute and test hierarchical F-statistics. *Mol. Ecol. Notes* **5**, 184 (2005).

35. K. Jordan *et al.*, A haplotype map of allohexaploid wheat reveals distinct patterns of selection on homoeologous genomes. *Genome Biol.* **16**, 48 (2015).

36. S. Wang *et al.*, Characterization of polyploid wheat genomic diversity using a high-density 90 000 single nucleotide polymorphism array. *Plant Biotechnol. J.* **12**, 787 (2014).

37. S. Salvi, O. Porfiri, S. Ceccarelli, Nazareno Strampelli, the 'Prophet' of the green revolution. *J. Agric. Sci.* **151**, 1 (2013).

38. H. Zhonghu;, S. Rajaram, Z. Y. Xin, G. Z. Huang, *A history of wheat breeding in China*. (CIMMYT, Mexico, 2001).

39. C. Hao *et al.*, The iSelect 9 K SNP analysis revealed polyploidization induced revolutionary changes and intense human selection causing strong haplotype blocks in wheat. *Sci. Rep.* **7**, 41247 (2017).

40. A. Betts, P. W. Jia, J. Dodson, The origins of wheat in China and potential pathways for its introduction: A review. *Quat. Int.* **348**, 158 (2014).

41. H. Tsujimoto, T. Yamada, T. Sasakuma, Pedigree of Common Wheat in East Asia Deduced from Distribution of the Gametocidal Inhibitor Gene (Igc1) and $\beta$-Amylase Isozymes. *Jppn. J; Breed.* **48**, 287 (1998).

42. Y. Zhou *et al.*, Uncovering the dispersion history, adaptive evolution and selection of wheat in China. *Plant Biotechnol. J.* **16**, 280 (2018).

43. A. L. Olmstead, P. W. Rhode, Adapting North American wheat production to climatic challenges, 1839–2009. *Proc. Natl. Acad. Sci. U.S.A.* **108**, 480 (2011).

44. Q. M. Paulsen, J. P. Shroyer, The early history of wheat improvement in the Great Plains. *Agron. J.* **100**, S70 (2008).

45. S. Rajaram, M. Van Ginkel, in *The World Wheat Book: A History of Wheat Breeding,* A. P. Bonjean, W. J. Angus, Eds. (Lavoisier, Paris, France, 2001), pp. 579-610.

46. A. P. Bonjean, W. J. Angus, *The World Wheat Book: A History of Wheat Breeding*. (Intercept, 2001).

47. R. K. Bacon, in *The World Wheat Book: A History of Wheat Breeding,* A. P. Bonjean, W. J. Angus, Eds. (Lavoisier, Paris, France, 2001), pp. 469-478.

48. C. J. Peterson, R. E. Allan, C. J. Peterson, in *The World Wheat Book: A History of Wheat Breeding,* A. P. Bonjean, W. J. Angus, Eds. (Lavoisier, Paris, France, 2001), pp. 407-430.

49. B. F. Carver, A. R. Klatt, E. G. Krenzer, in *The World Wheat Book: A History of Wheat Breeding,* A. P. Bonjean, W. J. Angus, Eds. (Lavoisier, Paris, France, 2001), pp. 445-468.

50. R. H. Busch, T. Rauch, in *The World Wheat Book: A History of Wheat Breeding,* A. P. Bonjean, W. J. Angus, Eds. (Lavoisier, Paris, France, 2001), pp. 431-444.

51. R. Joukhadar, H. D. Daetwyler, U. K. Bansal, A. R. Gendall, M. J. Hayden, Genetic Diversity, Population Structure and Ancestral Origin of Australian Wheat. *Front. Plant Sci.* **8**, 2115 (2017).
