## Supplementary figures and images for "Worldwide phylogeography and history of wheat genetic diversity"

### Supplementary file 2

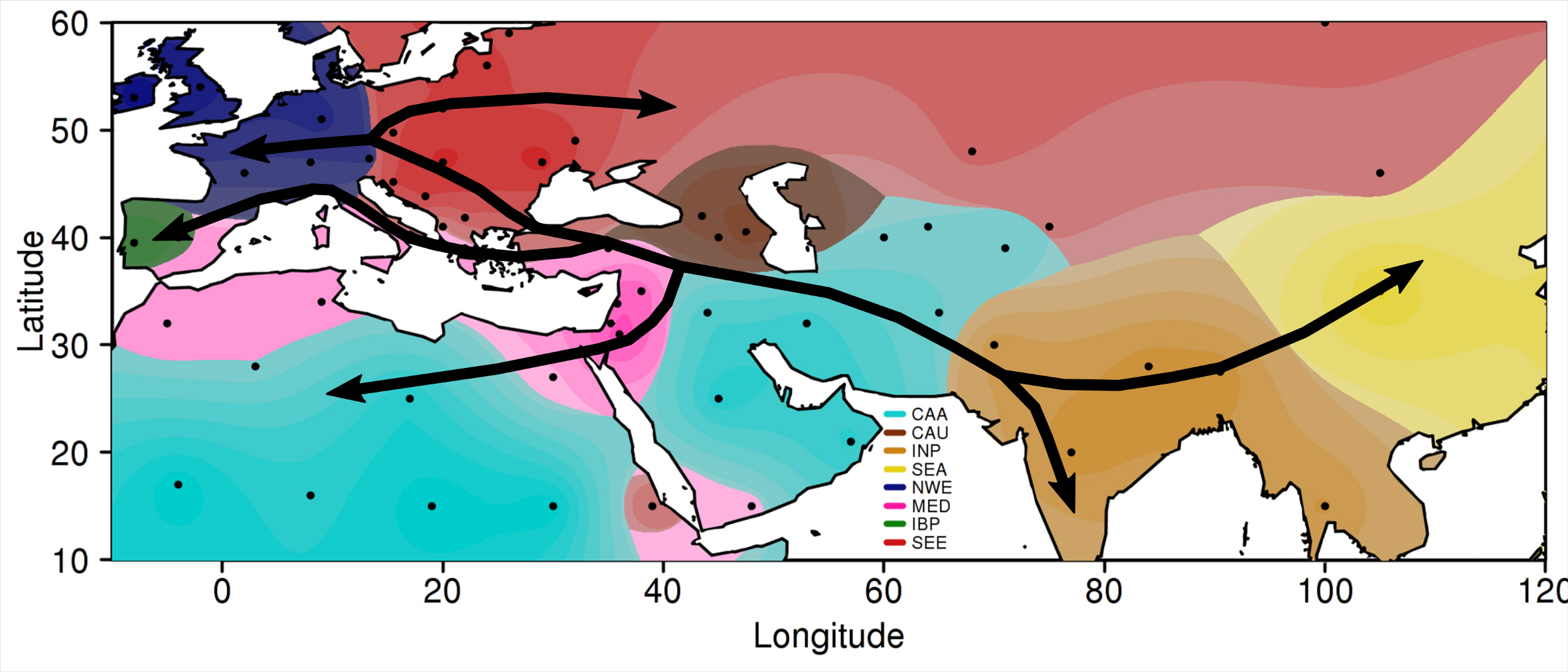

### Supplementary file 3

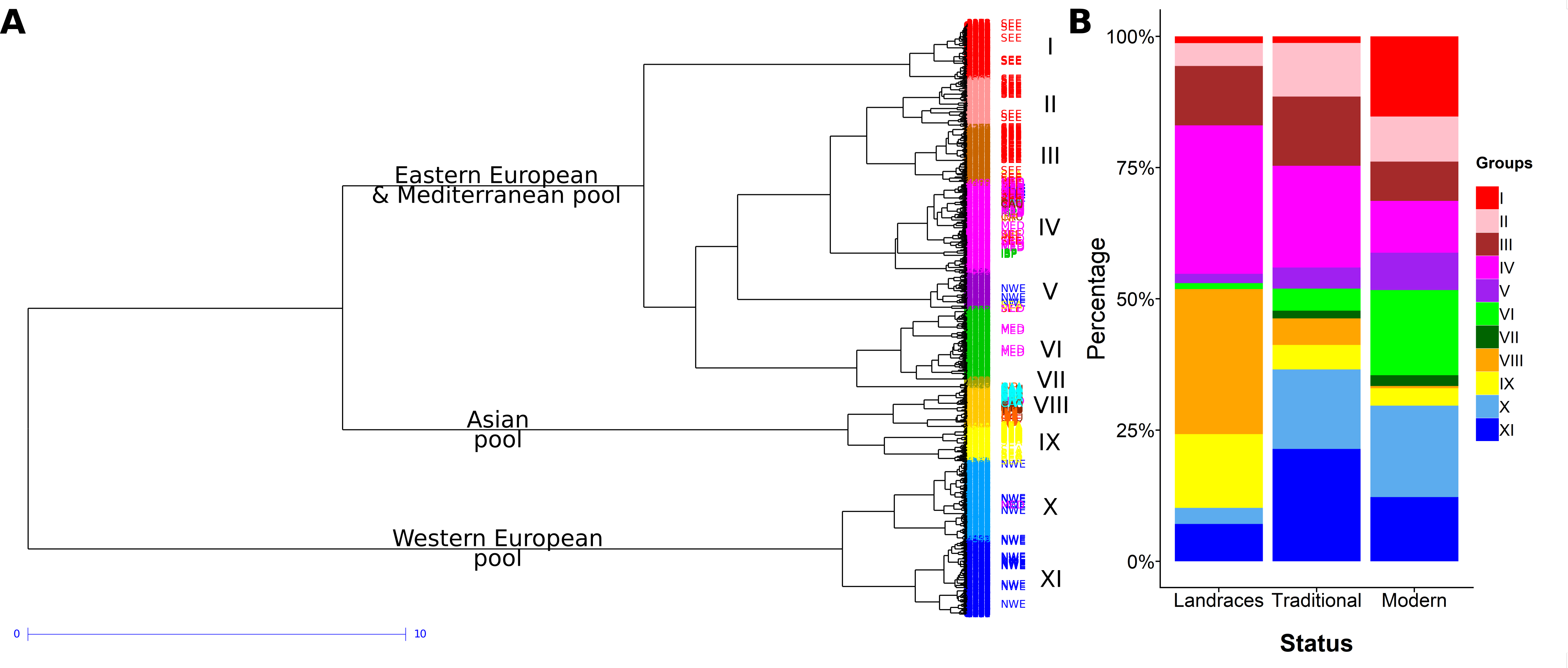

### Supplementary file 4

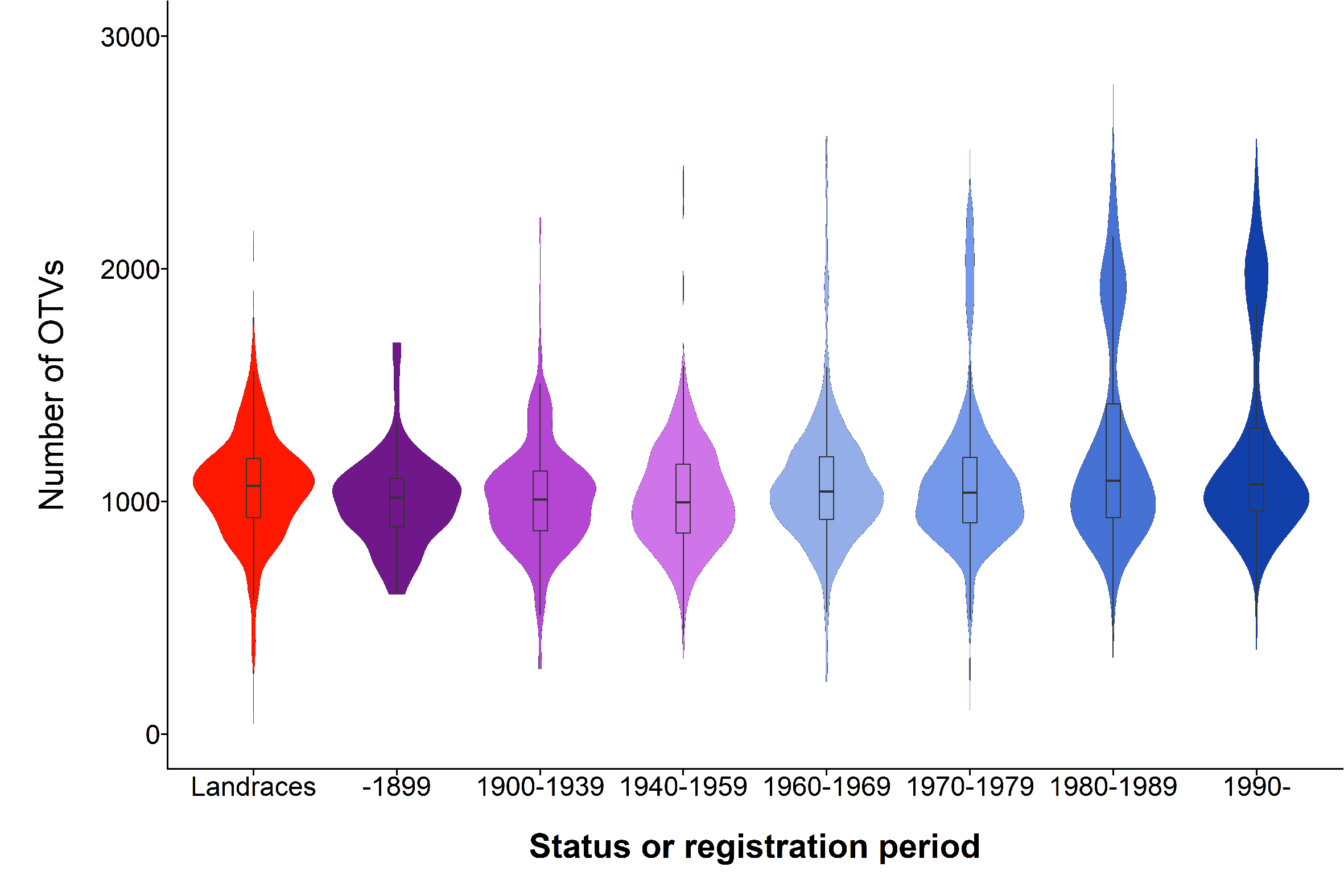
